# Reverse Watson-Crick purine-purine base pairs — the Sharp-turn motif and other structural consequences in functional RNAs

**DOI:** 10.1101/098723

**Authors:** Abhinav Mittal, Antarip Halder, Sohini Bhattacharya, Dhananjay Bhattacharyya, Abhijit Mitra

**Affiliations:** Center for Computational Natural Sciences and Bioinformatics (CCNSB), International Institute of Information Technology (IIIT-H), Gachibowli, Hyderabad 500032, India; Computational Science Division, Saha Institute of Nuclear Physics(SINP),1/AF, Bidhannagar, Kolkata 700064, India

**Keywords:** Sharp-turn motif, Noncanonical base pair, Reverse Watson-Crick base pair, Structural Bioinformatics, DFT, RNA

## Abstract

Identification of static and/or dynamic roles of different noncanonical base pairs is essential for a comprehensive understanding of the sequence-structure-function space of RNA. In this context, reverse Watson-Crick purine-purine base pairs (A:A, G:G&A:GW:W Trans) constitute an interesting class of noncanonical base pairs in RNA due to their characteristic C1′–C1′ distance (highest among all base pairing geometries) and parallel local strand orientation. Structural alignment of the RNA stretches containing these W:W Trans base pairs with their corresponding homologous sites in a non-redundant set of RNA crystal structures show that, as expected, these base pairs are associated with specific structural folds or functional roles. Detailed analysis of these contexts further revealed a bimodal distribution in the local backbone geometry parameters associated with these base pairs. One mode, populated by both A:A and G:G W:W Trans pairs, manifests itself as a characteristic backbone fold. We call this fold a ‘Sharp-turn’ motif. The other mode is exclusively associated with A:A W:W Trans pairs involved in mediating higher order interactions. The same trend is also observed in available solution NMR structures. We have also characterized the importance of recurrent hydrogen bonding interactions between adenine and guanine in W:W Trans geometry. Quantum chemical calculations performed at M05-2X/6-31++(2d,2p) level explain how the characteristic electronic properties of these W:W Trans base pairs facilitate their occurrence in such exclusive structural folds that are important for RNA functionalities.

## 1 Introduction

Non-coding functional RNAs have been found to be involved in a remarkable variety of tasks, ranging from enzymatic action to guiding chemical modifications in RNAs to regulating translation through temperature or metabolite sensing^1^. Recent progress in RNA-based therapeutics^2^ including RNA vaccination therapy^3^ and successful application of RNA based technologies in plant^4^ and animal agriculture^5^ have highlighted the importance of developing an overall understanding of the sequence-structure-function relationships in RNA. Despite numerous scientific investigations exploring the sequence-structure^6–9^ and structure-function^10–12^ spaces for RNA, a comprehensive picture similar to those for protein or DNA is yet to evolve^1, 13, 14^.

RNA achieves the remarkable structural complexity and fascinating functional diversity similar to proteins with only four different building blocks (nucleotides), instead of twenty physicochemically different amino acids as present in proteins. However, due to the presence of 2′-OH group of the RNA sugars, and the absence of methyl group at the C5-position of thymine, we observe diverse types of base pairing geometries in RNA apart from (a) the canonical G:C W:W Cis and A:U/T W:W Cis pairs^1^ present in double helical DNA and (b) some additional Hoogsteen type base pairs appearing in the telomeric regions of DNA. In the available RNA crystal structures, more than 85 types of such noncanonical base pairs have been detected which constitute about 30–40% of total instances of base pairing interactions in RNAs^15, 16^. In this context a reasonable hypothesis could be that different sets of noncanonical base pairs, because of their characteristic geometry and stability, may be involved in shaping RNA structures^17^ and dynamics^18^. A detailed investigation of sequence-structure-function relationships, which can be achieved by a combination of Quantum Mechanical (QM) and Bioinformatic studies, provides an important approach towards the validation of the hypothesis^19^. The utility of such a protocol has been demonstrated by Šponer and co-workers^20^ by showing that A:G W:W Cis base pairs occur at the ends of canonical helices with conservation and covariation pattern being determined by neighbourhood and tertiary interactions of guanine's pyramidal N2 amino group^20^. Similarly, consecutively occurring A:G H:S Trans base pairs are essential constituents of Kink-turn motifs^21,22^. Widely occurring Kink-turn motif introduces tight kink into RNA double helix and serves as binding site for number of proteins^23^. In case of GNRA tetraloops the first and last nucleotides engage in noncanonical A:G H:S Trans base pairing interaction^24^. Again, noncanonical U:A W:H Trans base pair is an essential component of the UA_handle submotif^25^. However, despite the rapid growth in the number of new RNA sequences and crystal structures, building a comprehensive structure-function annotation of other noncanonical base pairs is yet to be achieved.

Purine-purine reverse Watson-Crick (A:A W:W Trans, G:G W:W Trans and A:G W:W Trans) base pairs are known to occur at functionally important sites, such as, A14:A31 pair in Vitamin B12 aptamer^26^, G48:G71 pair in HIV-1 Rev peptide-RNA aptamer complex^27^, A1492:G530 pairing in the decoding center of large ribosomal subunit^28^, etc. They constitute an interesting class of noncanonical base pairs since the geometry of purine-purine W:W Trans base pairs are remarkable in two respects, (i) they have the maximum C1′-C1′ distance amongst all base pairing geometries and therefore causes the maximum stretch between the backbones, and (ii) in contrast to canonical Watson-Crick pairs, purine-purine Watson-Crick pairs interacting with their glycosidic bonds in *trans* orientation enforce a parallel local strand orientation. Therefore, in an otherwise double helical environment, where the local strand orientation is anti-parallel and composed of purine-pyrimidine base pairs, introduction of a purine-purine W:W Trans pair requires a special fold. Such exclusive structural folds are therefore expected to be associated with specific conformational dynamics or functional features of RNA. In this study we aim to identify such specific roles and understand their implications at the molecular level. Therefore, we have analyzed the available RNA crystal structures and performed a detailed bioinformatic survey of the structural neighborhood context, occurrence frequency, conservation and covariation patterns of the three purine-purine W:W Trans base pairs. We further have studied the ground state electronic properties and energetics of the structural components involving these base pairs using density functional theory (DFT) based approaches.

## 2 Tools and Methods

We selected 167 PDB files from non-redundant set of RNA crystal structures provided by HD-RNAs^29^ database for our analysis. HD-RNAs classifies RNA PDBs based on their functionality and source organism. Non-redundant dataset provided by HD-RNAs database contains representative structures decided upon after taking into account length, R-factor, resolution and sequence similarity. To exclude small synthetic RNA constructs, we selected only those PDBs which have resolution better than 3.5 Å and chain length greater than 30 nucleotides. We also have curated all the RNA X-ray crystal structures available in the PDB server on September, 2015 which has a resolution less than 3.5A (total 1873 structures). We will refer to this set of 1873 structures as ‘complete dataset’ and the former set of 167 structures as ‘non-redundant dataset’ in the following text. We have considered the solution NMR structures as well. Total 591 solution NMR structures containing RNA fragments, that were available in the PDB server on September, 2015 have been curated.

We have used precursor atom based algorithm called BPFIND^30^ to detect different base pairing interactions present in these 167 crystal structures. BPFIND detects base pairs involving at least two conventional hydrogen bonds (N-H…N, N-H…O, O-H…N, O-H…O, C-H…N and C-H…O type) formed between polar atoms of the bases or sugar O2′. In order to detect only ‘good’ base pairs we imposed following cutoffs criterion while using BPFIND — i) cutoff distance of 3.8 A between the acceptor and donor atoms ii) cutoff angle of 120.0 degree for checking planarity of precursor atoms as well as linearity of hydrogen bonds and iii) cutoff ‘E-value’ of 1.8 to signify the overall distortion and maintain a good base pairing geometry. Details of the occurrence frequencies of different base pairing geometries in the non-redundant set of RNA crystal structures are available in RNABP COGEST database^31^. However, due to cutoff based base pairing criteria, a number of physically relevant base pair instances remain undetected.

To identify these undetected base pairs and to check for conservation and covariation pattern of base pairs across the species, structural alignment study guided by results from BPFIND has been performed. Structural alignment is necessary for this purpose, since, homologous RNAs from different organisms have slightly different sequence and structure and consequently have different residue numbering. We have used SETTER^32–35^ algorithm for structural alignment of RNAs. For the purpose of 3D structural alignment, SETTER (SEcondary sTructure-based TERtiary Structure Similarity Algorithm) divides an RNA structure into a set of non-overlapping structural elements called generalized secondary structure units (GSSUs). Structural alignment of homologous RNA chains has been followed by structural alignment of smaller homologous neighbourhood fragments. This approach provides us with a robust platform to study conservation and covariation patterns of non-bonding interactions including base pairing interactions and other structural features across species. VMD^36^ and PyMol^37^ are used for visualization and image rendering. NUPARM^38^ package is used for calculating intra base pair paramters.

In order to study the ground state electronic properties of different nucleobases and different structural fragments containing them, first we have modeled them appropriately. Modeling has been done by extracting the coordinates of different bases, base pairs, base triples and quartets from the crystal structures followed by adding hydrogen atoms at appropriate positions. For our calculations we have considered three models of a nucleobase depending on the functional group that is present at N9 of purines and N1 of pyrimidines, *e.g.,* (i) nucleobases with only hydrogen at N9/N1 position, (ii) nucleobases with a methyl group at N9/N1 position and (iii) nucleobases with the complete sugar moiety at N9/N1 position. For the third case we have removed the phosphate group and added a methyl group at O3′ position. In the crystal structures we have identified some examples where one of the sugar moiety of a base pair is present in unusual *syn* orientation. We have considered those examples also. However only the models with methyl substitution have been reported in the main text and rest of the models have been discussed in the supporting information. Crystal geometry of these molecular fragments have been optimized to obtain the minimum energy structure using Density Functional Theory (DFT). For geometry optimization and subsequent interaction energy calculations, we have selected Truhlar's M05-2X ^39^ functional and the 6-31G++(2d,2p) basis set^40^. M05-2X is a hybrid meta GGA functional with 56% Hartree-Fock exchange component and is known to perform remarkably well for noncovalently bonded molecules of biological relevance,^41^ including nucleobases^42–49^. 6-31G++(2d,2p) is a split valance double ζ type basis set, with two sets of d type polarization function for all non-hydrogen atoms and two sets of p type polarization function for hydrogen atoms, and also include s-p diffused orbitals for non-hydrogen atoms, was used for ground state geometry optimization. Two types of geometry optimization have been performed, (i) H*_opt_* (where all the non-hydrogen atoms are kept fixed at their crystal structure coordinates and only the hydrogen atoms are allowed to change its position) and (ii) F_*opt*_ (where no constraints are applied on the atoms). H_*opt*_ geometries are therefore crystal geometries with hydrogen atoms placed at minimum energy positions. We have performed a Hessian calculation for all these optimized geometries to confirm that, the corresponding Hessian matrices have only real eigenvalues. We have further calculated the interaction energies of the base pairs (n=2)/triples (n=3)/quartets (n=4) as 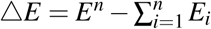, where E^*n*^ in the electronic energy of the ground state optimized geometry of the complex and E, is the electronic energy of the ground state optimized geometry of the individual monomer i. We have corrected the interaction energies ( ΔE) for Basis Set Superposition Error (BSSE) and Zero Point Vibrational Energies (ZPVE) and the final corrected interaction energy is denoted as E^*int*^. BSSE calculations have been performed using Boys's counterpoise method^50^. Note that, although implementing BSSE corrections via counterpoise method has recently faced some serious criticism^51, 52^, it is well accepted for DFT based approaches in base pairing or equivalent systems^53,54^. For all the QM calculations and molecular editing we have used Gaussian 09 (Revision C.01)^55^ and GaussView^56^ packages respectively.

## 3 Results and Discussions

All the occurrences of purine-purine reverse Watson-Crick (W:W Trans) base pairs in the HD-RNAS dataset have been archived in RNABP COGEST database, which reports 139 occurrences of A:A W:W Trans, 17 occurrences of G:G W:W Trans and 6 occurrences of A:G W:W Trans.^2^ Some of these base pairs are found to participate in base triple formation, where the free sugar and Hoogsteen edges of the W:W Trans pair interact with a third base to form a second planar base pair with two hydrogen bonds. Such triples are also reported in ‘Non-canonical RNA Base Pair Database’ (http://www.saha.ac.in/biop/www/db/local/BP/rnabasepair.html), and are accessible through RNABP COGEST. Again, free edges of the W:W Trans pair may interact with two other bases simultaneously and form a base quartet. All such instances detected in HDRNAS are listed in QUARNA server (http://quarna.iiit.ac.in/).

### 3.1 Intrinsic stability of isolated A:A, G:G and A:G W:W Trans pairs

We have obtained the gas phase ground state optimized geometries of the A:A W:W Trans, G:G W:W Trans and A:G W:W Trans pairs at M05-2X/6-31G++(2d,2p) level of theory and have calculated their BSSE and ZPVE corrected interaction energies (E^*int*^) at the same level of theory. From the NUPARM parameters reported in Figure 1 we note that, optimized geometries of A:A W:W Trans and G:G W:W Trans base pairs are significantly planar and stabilized by 2 N-H…N and 2 N-H…O type bonds, respectively. Although we did not impose any symmetry based restriction during geometry optimizations, the optimized structures show nearly C2 symmetry about an axis passing through the center and perpendicular to the base pair plane. We also observe that, though the amino group of unpaired adenine has an intrinsic pyramidal geometry, as is expected due to a partial sp^3^ hybridization at N6, in the optimized geometry of A:A W:W Trans, as reported earlier,^57,58^ the amino groups are found to be in planar sp^2^ arrangement due to their participation in in-plane hydrogen bonding interaction (Figure S2 of supporting information).

**Figure 1.**
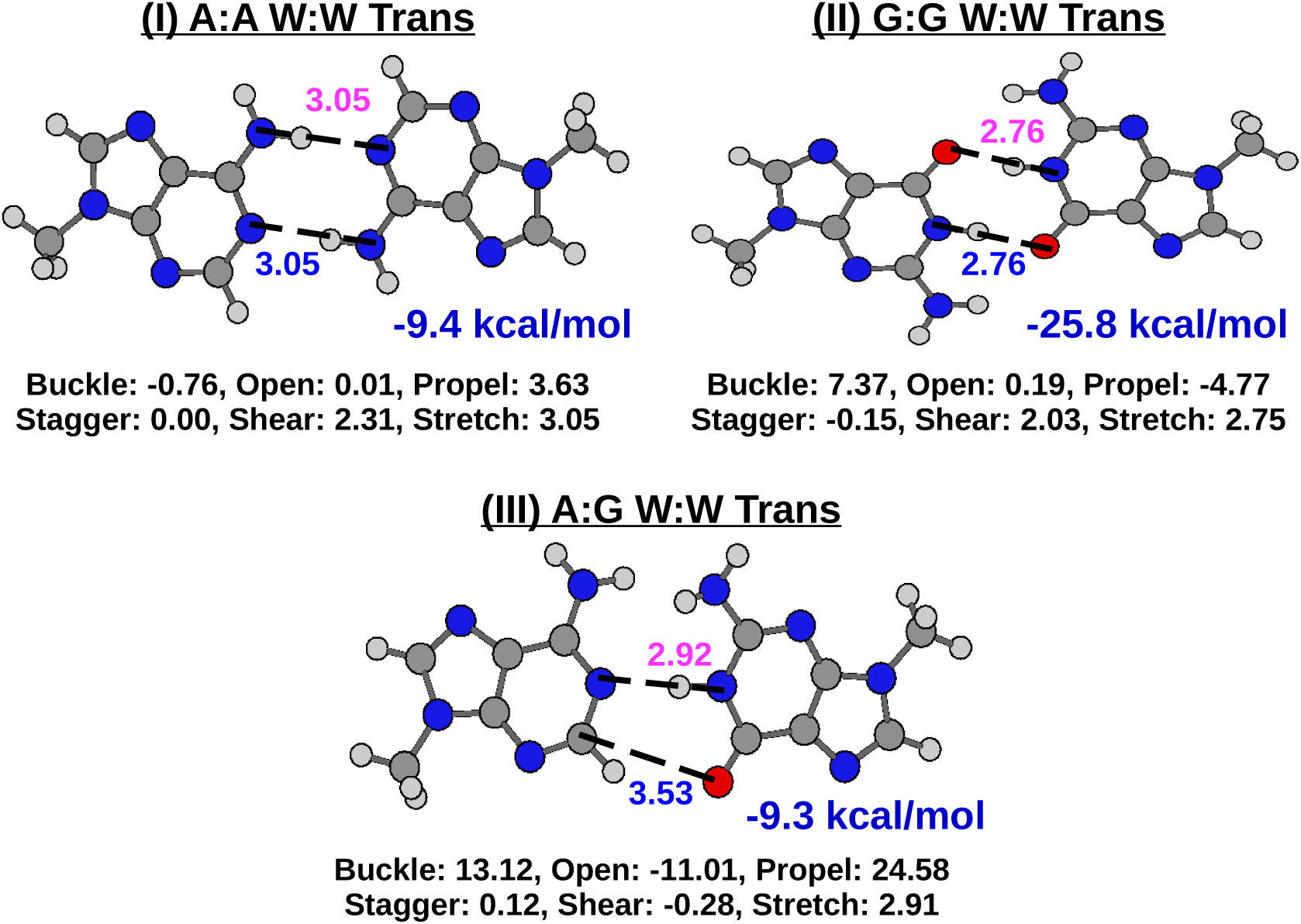
Optimized geometries and interaction energies (E^*int*^) of A:A, G:G and A:G W:W Trans pairs at M05-2X/6-31G++(2d,2p) level of theory. Sugar moieties at N9 positions have been substituted by methyl groups. Inter base pair rotational (Buckle, Open, Propeller) and translational (Stagger, Shear, Stretch) geometric parameters calculated by NUPARM package have been reported. Donor-acceptor distances for all the inter base hydrogen bonds are given in Å. Similar results for different substitutions at N9 positions have been reported at Figure S1 and Table S1 of supporting information.

Interestingly, in contrast with that of the homopurine pairs, the optimized geometry of A:G W:W Trans is non-planar, and is characterized by high Buckle and Propeller values, with one N-H…N type and a C-H…O type hydrogen bonds (Figure 1, S1 and Table S1). Due to such inherent non-planar geometry of A:G W:W Trans, it has a shorter *C*1′ — *C*1′ distance (12.92 Å) compared to A:A (13.75 Å) and G:G (13.12 Å) W:W Trans pairs.^3^ Note that, A:A and G:G W:W Trans base pairs have been classified as isosteric by Leontis and co-workers in their classical work on isostericity of base pairs^59^. However, A:G W:W Trans remained uncharacterized because it has not been identified as a base pair^60^. Using the BPFIND algorithm, we reported,^31^ 6 instances of A:G W:W Trans pairing interactions in the non-redundant set of RNA crystal structures. Our QM calculations show that, except in one case, these recurrent hydrogen bonded interactions are significantly stable, as supported by the small RMSD (<0.5Å) between their respective H_*opt*_ and F_*opt*_ geometries (Table S11 and Figure S7 of Supporting information). We have also found that A:G W:W Trans, if considered as a base pair, is ‘near-isosteric’ with the other two purine-purine W:W Trans base pairs. Details of the IsoDiscrepancy Index (IDI)^16^ calculation of A:G W:W Trans with respect to A:A W:W Trans and G:G W:W Trans are given in Supporting Information.

All the three purine-purine reverse Watson-Crick base pairs (A:A, G:G and A:G) are composed of two conventional hydrogen bonds: two N-H…O bonds in G:G W:W Trans, two N-H…N bonds in A:A W:W Trans and one N-H…N and one C-H…O bonds in A:G W:W Trans. Among them, the N-H…O type hydrogen bonds present in G:G W:W Trans pair are remarkably stronger than other inter-base hydrogen bonds, as inferred by the high red shift (>475cm^−1^) in the symmetric stretching frequencies of the N-H bond (Table 1). As a consequence of such strong inter-base hydrogen bond formation, interaction energy of G:G W:W Trans pair is nearly 2.5 times higher than those of A:A and A:G W:W Trans pairs (Figure 1 and Table S2). In A:G W:W Trans, although the G(N1-H)…(N1)A hydrogen bond (colored in magenta in Figure 1) is stronger (red shift = 457.5 cm^−1^) than the N-H…N bonds of A:A W:W Trans, the second hydrogen bond (A(C2-H)…(O6)G) is significantly weaker with a small red shift of 26.8 cm^−1^. We also observe that, for the base pairs modeled with sugar at the N9 position, as expected, the *anti* orientation of the sugar results in higher interaction energies, specially for A:G W:W Trans (Table S2 of Supporting Information). Interestingly, the occurrence frequencies of these purine-purine base pairs do not correlate with the intrinsic strengths of their interactions. This is in contrast with the trend we observe in case of canonical base pairs where, although G:C W:W Cis and A:U W:W Cis are isosteric to each other, occurrence frequency of stronger G:C W:W Cis base-pair is approximately 4 times that of weaker A:U W:W Cis base-pair.

**Table 1.** Shift in the frequency of vibration (symmetric stretch) of the hydrogen bond donors due to formation of base pairs (with methyl at N9 position), triples and quartets. *v*_0_ and *v* represent the frequency before and after the inter base hydrogen bond formation, respectively (in cm^−1^ unit). *I*_0_ and *I* represent the IR intensity before and after the inter base hydrogen bond formation, respectively (in KM/mole unit). For base triples and quartets we have reported the hydrogen bonds of only the A:A W:W Trans pair. Hydrogen bond donor-acceptor distances of the H-bond I and H-bond II have been colored as magenta and blue, respectively, in Figures 1, 6 and 8. Table S3 of supporting information lists same parameters for base pairs with different N9 substitutions.

| System | H-bond I (magenta) |  | H-bond II (blue) |  |
| --- | --- | --- | --- | --- |
| | $\nu_0 - \nu$ | $\frac{I}{I_0}$ | $\nu_0 - \nu$ | $\frac{I}{I_0}$ |
| **Base Pairs** |  |  |  |  |
| A:A W:W Trans | 261.8 | 25.1 | 261.8 | 25.1 |
| G:G W:W Trans | 477.3 | 58.2 | 477.3 | 58.2 |
| A:G W:W Trans | 457.5 | 25.4 | 26.8 | 2.9 |
| **Base Triples** |  |  |  |  |
| A:A-U | 248.0 | 21.4 | 277.0 | 0.6 |
| A:A-G | 290.1 | 4.7 | 234.6 | 18.3 |
| A:A-C(+) | 501.9 | 15.2 | 153.9 | 8.1 |
| A:A(+)-C | 590.3 | 13.1 | 122.7 | 6.9 |
| **Base Quartets** |  |  |  |  |
| A:A-U-U | 239.6 | 18.7 | 239.6 | 18.7 |
| A:A-C(+)-U | 528.9 | 17.7 | 171.6 | 5.4 |
| A:A(+)-C-U | 628.8 | 14.0 | 174.5 | 5.3 |
| A:A-G-U | 289.0 | 5.5 | 227.0 | 16.2 |
| A:A-G-G | 229.1 | 3.2 | 204.4 | 23.2 |

Our QM calculations suggest that, the intrinsic stabilities and geometries of the three purine-purine W:W Trans base pairs are significantly different from each other. Even the isosteric A:A and G:G pairs are quite different in terms of their inter-base hydrogen bonds and physicochemical surface. In the following section we will explore how do the characteristic geometries and electronic properties of these W:W Trans base pairs determine their occurrences at the sites with specific functional roles in RNA.

### 3.2 Bimodal classification of all the occurrence contexts on the basis of local backbone geometry

Note that, A:A and G:G W:W Trans pairs being isosteric to each other and near isosteric to A:G W:W Trans, are expected to occur at similar structural folds defined by the local backbone geometry. We have characterized the local geometry of a base pairing interaction (between i^*th*^ and j^*th*^ residues) by the following two parameters — (i) their *C*1′(*i*) − *C*1′(*j*) distance and (ii) the combination of the backbone angles *ρ_i_* and *ρ_j_*, for i^*th*^ and j^*th*^ residues, respectively. As described in Figure 2a, *ρ_n_* is the angle formed between the three phosphate atoms of three successive residues, i.e., ∠ *P*(*n − 1*) − *P*(*n*) — *P*(*n* + 1). A base pair involving i^*th*^ and j^*th*^ residues will therefore have two backbone angles, *ρ_i_* for i^*th*^ residue and *ρ_j_* for j^*th*^ residue. We will use the term *ρ_min_* to indicate the smaller of the two values *ρ_i_* and *ρ_j_*. As a reference for comparison, we have considered the average values of *ρ_i_* (*ρ̅*_*i*_) for the standard A and B type nucleic acid helices: for A-DNA and A-RNA *ρ̅*_*i*_ = 151.5°, for B-DNA *ρ̅*_*i*_ = 147.4°. For a GNRA teraloop, *ρ_i_* values corresponding to the terminal nucleotides (G and A) are of the same order as B-DNA and are usually higher compared to the central nucleotides (N and R) as reported in Supporting Information.

**Figure 2.**
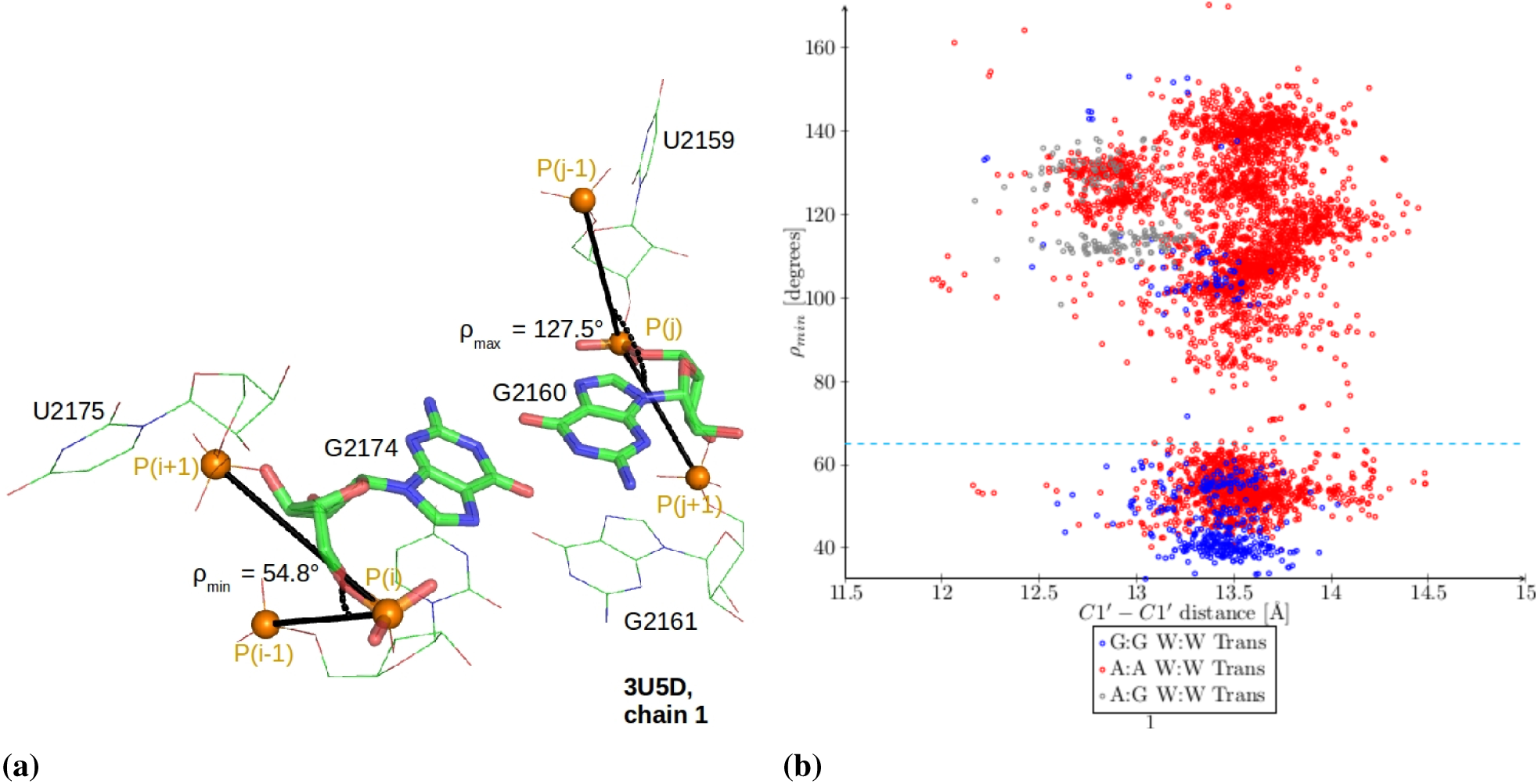
(a) Definition of *ρ_min_* and *ρ_max_* angles are shown for a G:G W:W Trans base pair. (b) *ρ_min_* angle vs. *C*1′ — *C*1′ distance have been plotted for purine-purine Watson-Crick Trans base pairs detected by BPFIND in complete dataset. The dotted blue line represents the *ρ_min_* = 65° value.

Analyzing the complete set of RNA crystal structures we have observed that, (a) among all possible base pairing geometries, the value of *C*1′(*i*) — *C*1′(*j*) distance is the maximum for purine-purine W:W base pairs^4^ and (b) a significant amount of purine-purine W:W Trans base pairs occur with either *ρ_i_* or *ρ_j_* (not both) lying within the range 45 to 65 degrees implying a sharp turn on one of the strands. We explored the phase space defined by the minimum of *ρ_i_* and *ρ_j_* (*ρ_min_*) and the *C*1′(*i*) — *C*1′(*j*) distance for all the purine-purine W:W Trans base pairs (*i*, *j*) detected in the complete set of RNA crystal structures, and observed that the occurrences of purine-purine W:W Trans base pairs are clustered into two distinct regions (Figure 2b). Interestingly, barring a few exceptions occurrences of G:G W:W Trans base pairs are limited to the region defined by small *ρ_min_* values (<65°) and occurrences of A:G W:W Trans base pairs are limited to the region defined by large *ρ_min_* values (>65°). In contrast, A:A W:W Trans base pairs are distributed over both the regions. The same clustering can also be visualized in the plot of *ρ_min_* angles with respect to *ρ_max_* angles (Figure S3 in supporting information).

### 3.3 Structural context common to A:A W:W Trans and G:G W:W Trans — the Sharp-turn motif

The discussion above is based on good base pairs defined by BPFIND software. For a more comprehensive context analysis we have also considered other instances with similar geometries but with a more relaxed base pairing criterion as discussed in the methods section. This increased the the count of A:A W:W Trans base pairs with affecting those of G:G and A:G W:W Trans base pairs. We have also studied the contexts of the A:A and G:G W:W Trans base pairs in terms of their (i) influence on backbone geometry, (ii) other non-bonding interactions and (iii) identity of neighboring residues. Our analysis shows that, in the non-redundant set of RNA crystal structures, a significant number of A:A W:W Trans base pair instances (40 out of 155) and almost all of G:G W:W Trans base pair instances (11 out of 17) have characteristic small *ρ_min_* values (*ρ_min_* < 65° region in Figure 2). We observe that, these instances are immediately preceded or succeeded by a canonical base pair. The arrangement imposes a sharp turn in the backbone of one of the two strands as shown in Figure 3 A and B, and schematically described in Figure S4. The sharp turn in backbone is required to accommodate a canonical base pair which has anti-parallel local strand orientation and 10.7 Å C1′-C1′ distance, adjacent to a A:A W:W Trans or G:G W:W Trans base pair which has parallel local strand orientation and 13.6 Å C1′-C1′ distance. It is noteworthy that, the sharp turn brings negatively charged phosphate groups in close vicinity (Figure 3). To balance the resulting electrostatic repulsions, metal cations in association with water molecules and charged amino acid residues are found to interact with the Sharp-turn motifs. We have identified a number of such cases, specially in 23S rRNA of different species. One example from the 23S rRNA of *H. marismortui* has been shown in Figure 3 C. Other examples from 23S rRNA of *E. coli, T. thermophilus* and *H. marismortui* have been elaborated in Figure S7 of Supporting Information.

**Figure 3.**
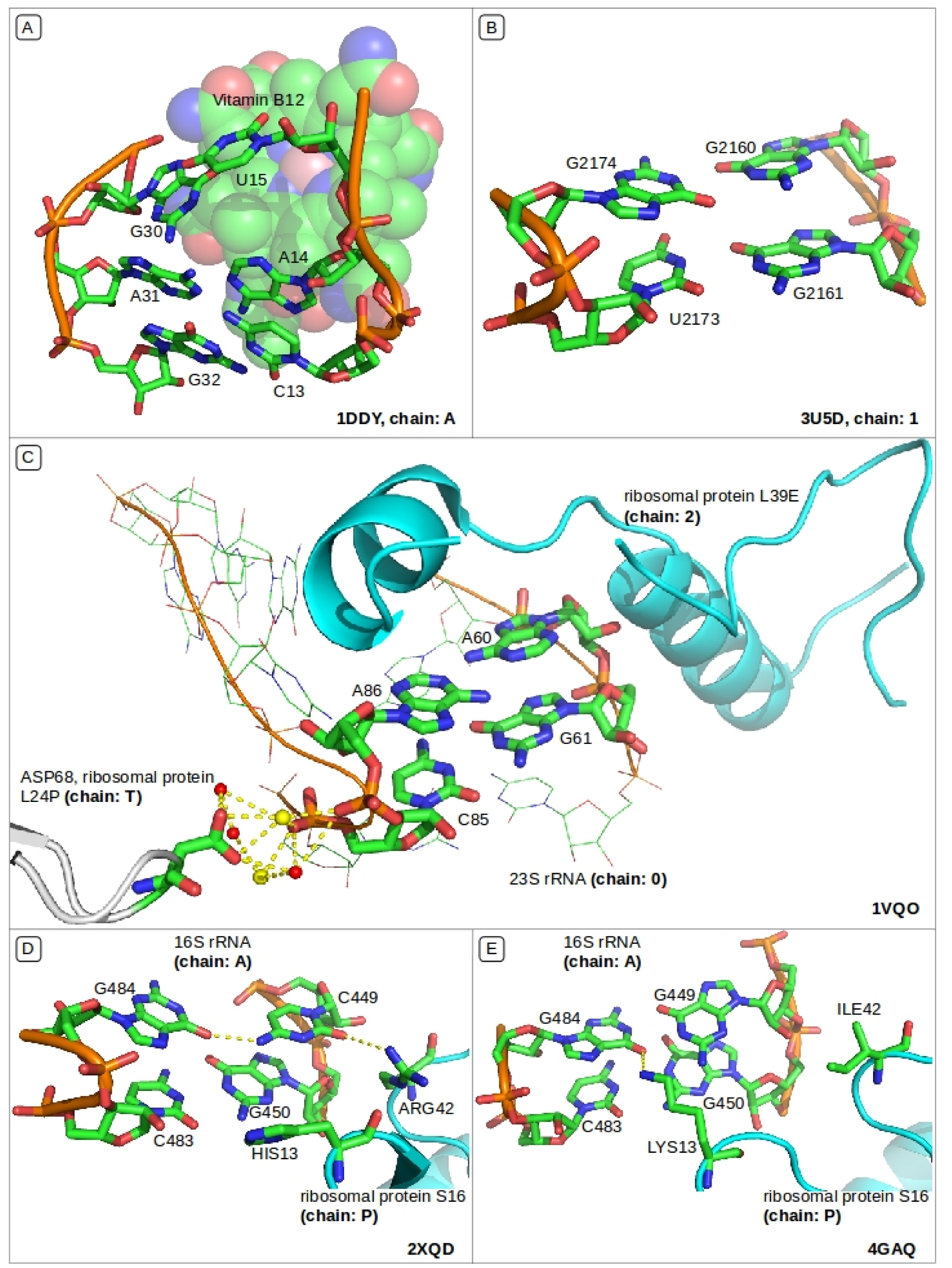
**(A)** Sharp-turn motif composed of occurrences of G:C W:W Cis and A:A W:W Trans base pairs and forming binding pocket for vitamin B12; **(B)** Sharp-turn consisting of consecutively occurring of G:G W:W Trans and G:U W:W Cis; **(C)** Sharp-turn interacting with ribosomal protein and ions (strontium ions in yellow and waters in red); **(D)** and **(E)** are homologous occurrences of Sharp-turn interacting with ribosomal protein S16 in 16S rRNA from *T. thermophilus* and *E. coli,* respectively. The bases of Sharp-turn in 16S rRNA and amino acids of protein S16 have co-evolved.

In the subsequent discussion we will refer to this particular structural arrangement as ‘Sharp-turn’. We have listed all the occurrences of Sharp-turns in Table S7 of supporting information, which shows that, in the non-redundant set of RNA crystal structures there are a total of 36 PDB files (including 23S rRNA, 16S rRNA and Vitamin B12 RNA aptamer) which contain the Sharp-turn composed by A:A or G:G W:W Trans pairs. Interestingly, we have identified that, Sharp-turn occurs as a part of important structural motifs such as, internal loop, kissing loop, junction loop and pseudoknot (Table S7). It may be noted that, similar sharp turn in the backbone can also be characterized by conventional backbone torsion angles of RNA, such as η and *θ*. However, we consider that use of *ρ_n_* angle is more straight forward, because otherwise it requires both the η of the i ^*th*^ and *θ* of the i-1^*th*^ residues, respectively to characterize a Sharp-turn (Figure S10, Table S17 of Supporting Information). As shown in Table S17, for a Sharp-turn the value of η of the i^*th*^ residue lies between 43.0° to 76.0° and the value of *θ* of the i-1^*th*^ residue lies between −50.0° to −24.0°. It may be noted that both η and *θ* values are around 180° in regular double helical structures consisting of Watson-Crick base pairs.

#### 3.3.1 Composition and conservation of Sharp-turn

We have identified 78 instances of Sharp-turn which are further categorized into 9 different contexts — 6 in 23S rRNA, 2 in 16S rRNA, 1 in 25S rRNA and 1 in vitamin B12 aptamer (Table S7). Structural alignment of 23S and 16S rRNAs available in the data set reveals that, the Sharp-turn motif is conserved across species, however its composition contains a number of variations. These variations include, (i) substitution of a purine base of the A:A or G:G W:W Trans pair by a pyrimidine base (e.g. context 1 and context 5), (ii) substitution of both the purine bases by pyrimidine bases (e.g., context 2), (iii) alteration of purine-purine base pairing geometry (e.g. context 8), (iv) difference in the type of canonical base pair that occurs adjacent to the W:W Trans pair (e.g., context 1). We have identified that the A:A and G:G W:W Trans pairs, which are involved in mediating the Sharp-turn, are also involved in non-covalent interactions with other residues. Such interactions remain unchanged even if one or both of the purine bases get substituted by pyrimidine bases. One such example is observed in 23S rRNA of *Haloarcula marismortui* (PDB: 1VQO), where both the adenine residues got substituted (by C768 and U893 residues, respectively), but exhibit similar interaction pattern with C162 as observed in the other homologous sites (e.g., A678, A800 and C192 of *T. thermophilus* 23S rRNA), see Figure 4.C and D. Another interesting example is observed in the context 4, where the local backbone geometry of Sharp-turn is not conserved, in 1VQO, as shown in Figure 4 A and B. The RNA chain takes a longer path to compensate the lack of Sharp-turn motif and maintains the overall folding pattern. However, in this case also interaction of the G1025:G1139 W:W Trans base pair with the A1143 residue observed in 23S rRNA from *E. coli* (Figure 4 A) remains conserved in 1VQO structure where A1242:U1122 W:W Cis base pair interacts with A1247 (Figure 4 B). Figure 4 E and F describe the examples of context 1, where a Sharp-turn motif composed of A:A W:W Trans is present in the 23S rRNA of *E. coli, H. marismortui* and *D. radiodurans,* and is associated with hydrogen bonding interactions with the neighboring uracil residue (U68 in 3R8S). In *T. thermophilus,* interaction with the neighboring uracil remains conserved even when one of the adenines of the A:A W:W Trans pair gets substituted by a uracil.

**Figure 4.**
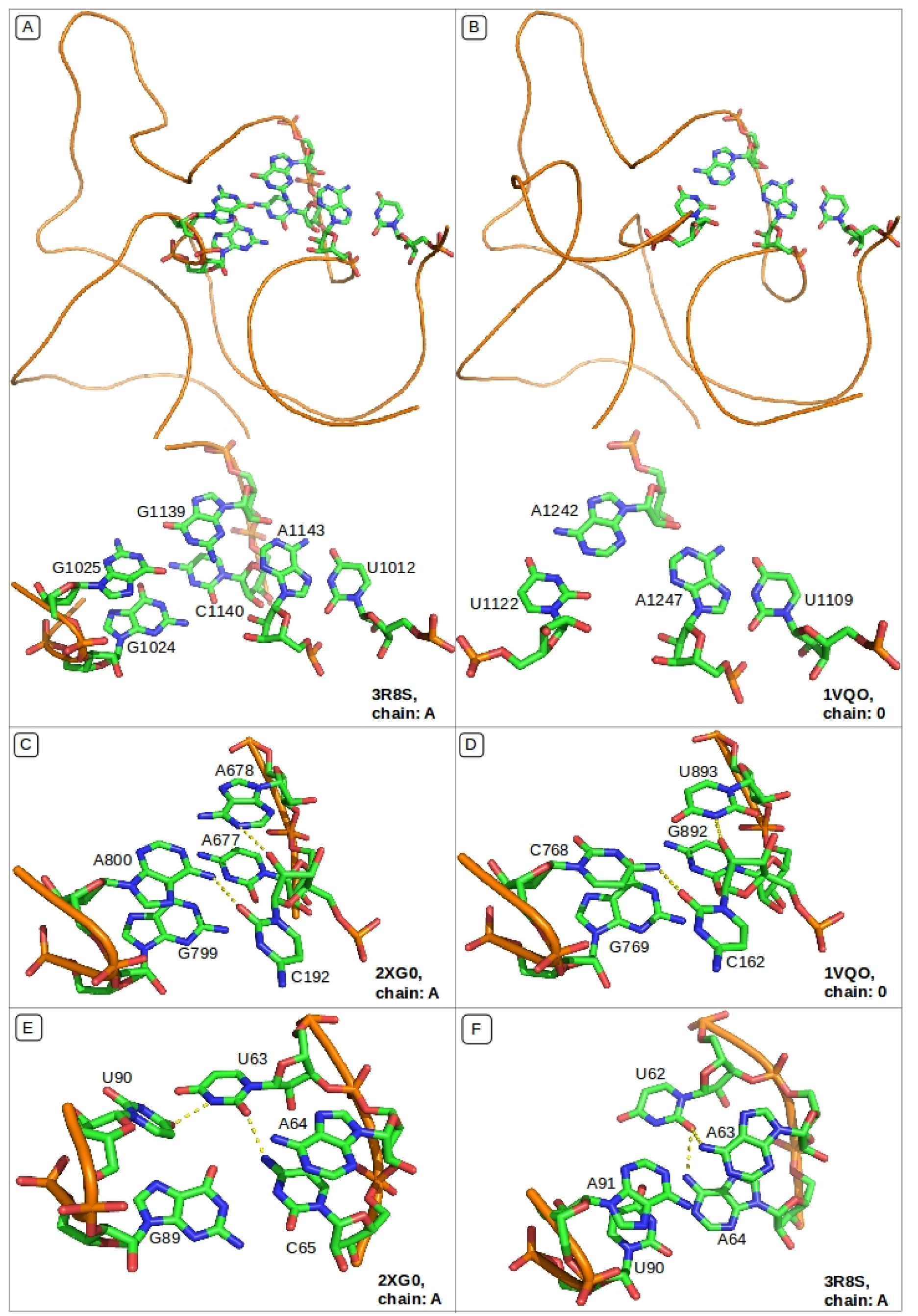
**(A)** and **(B)** Lack of conservation of Sharp-turn motif in 23S rRNA from *H. marismortui* (1VQO) and conservation of the tertiary interactions with adenine; **(C)** and **(D)** Conservation of backbone fold and Hydrogen bond interactions of Sharp-turn mediated by A:A W:W Trans and C/U bases; **(E)** and **(F)** Conservation of backbone fold and interactions with nearby residue despite variation in composition of Sharp-turn

#### 3.3.2 Biological Significance of Sharp-turn

Our analysis reveals occurrences of Sharp-turn motifs at structurally important sites of functional RNAs. Here we discuss three interesting examples.

##### Vitamin B12 Aptamer

Cobalamin riboswitch was the first riboswitch shown to directly interact with a metabolite in absence of proteins^61^. These riboswitches have been found to regulate B12 biosynthesis in bacteria^62^ and are widespread in prokaryotes^63,64^. In synthetically SELEX-generated vitamin B12 aptamer 1DDY, an important component of the binding pocket is a Sharp-turn consisting of A:A W:W Trans base pair followed by G:C W:W Cis canonical pair (Figure 3 A, context 7 in Table S7). Binding of vitamin B12 is stabilized by complementary packing of hydrophobic surfaces, hydrogen bonding and dipolar interactions^26^. Interestingly, in the binding of naturally occurring cobalamin aptamers, the molecular details of recognition were found to be quite different and does not involve the Sharp-turn mediated architecture^65^. Instead, a whole new set of interactions are brought into play.

##### Interaction with ribosomal proteins I

We have detected another Sharp-turn (composed of A:A W:W Trans and G:C W:W Cis) that provides the interface for interaction between two ribosomal proteins, L39E and L24P, and 23S rRNA of *H. marismortui* (1VQO). However the interaction pattern for L39E and L24P are quite different from from each other. For L39E, one of its two alpha helices directly interacts with this Sharp-turn. Whereas, L24P interacts indirectly with the Sharp-turn via a network of non-covalent interactions formed by (i) the aspartic acid residue of the protein (residue no. 68, chain T), (ii) the negatively charged phosphates (in the U-shaped backbone of A-C base-step) and (iii) strontium ions and water molecules present in the crystal (Figure 3 C).

##### Interaction with ribosomal proteins II

In 30S ribosomal subunit of *E. coli*, we observe a Sharp-turn consisting of G:G W:W Trans and G:C W:W Cis, where the G:G W:W Trans base pair is composed of G450 and G483 residues (Figure 3 E). G450 residue further interacts with amino group of LYS13 residue of ribosomal protein S16. S16 also has a non-polar and noninteracting ILE42 residue close to the Sharp-turn. At homologous location in 30S ribosomal subunit of *T. thermophilus,* we observe a variant of the same Sharp-turn where the G483 residue has been replaced by a cytidine residue (C483) (Figure 3 D). Note that, only N4 amino group of C483 interacts with O6 of G450 keeping the C1′-C1′ distance (13.3 Å) similar to that of a G:G W:W Trans base pair. Such change in the composition of Sharp-turn motif affects the RNA-protein interaction network. Hence we observe that, in *T. thermophilus* HIS13 and ARG42 are present instead of LYS13 and ILE42, of *E. coli.* Interestingly in *T. thermophilus* O2 of C483 interacts with amino group of ARG42 and the non-polar aromatic HIS13 remains noninteracting. Our observations indicate that, 16S rRNA and ribosomal protein S16 co-evolved, preserving the binding interaction between the Sharp-turn of RNA and the protein (Figure 3 D, E).

### 3.4 Structural context specific to A:A W:W Trans

As shown in Figure 2b, the population of the region with high *ρ_min_* value, is dominated by A:A W:W Trans base pairs. Survey of structural context of these A:A W:W Trans base pair instances reveals that majority of them occur as part of higher order interactions, such as base triples and quartets. One or both of the free Hoogsteen edges of these A:A W:W Trans pairs further pair with sugar edge of guanine or Watson-Crick edge of uracil or cytosine in *trans* fashion. Table S8 and Table S9 list all such higher order interactions involving A:A W:W Trans as found in the non-redundant dataset. Our context analysis reveals that, these triples and quartets either mediate loop-loop interactions or are part of junction loops (Figure 5, 7 and Table S8, S9). Such long-range tertiary interactions and junction loops are crucial to the maintenance of structure and function in RNAs^66^. In the next section we discuss the energetics of such higher order interactions along with their importance in maintaining the tertiary structure of RNA.

**Figure 5.**
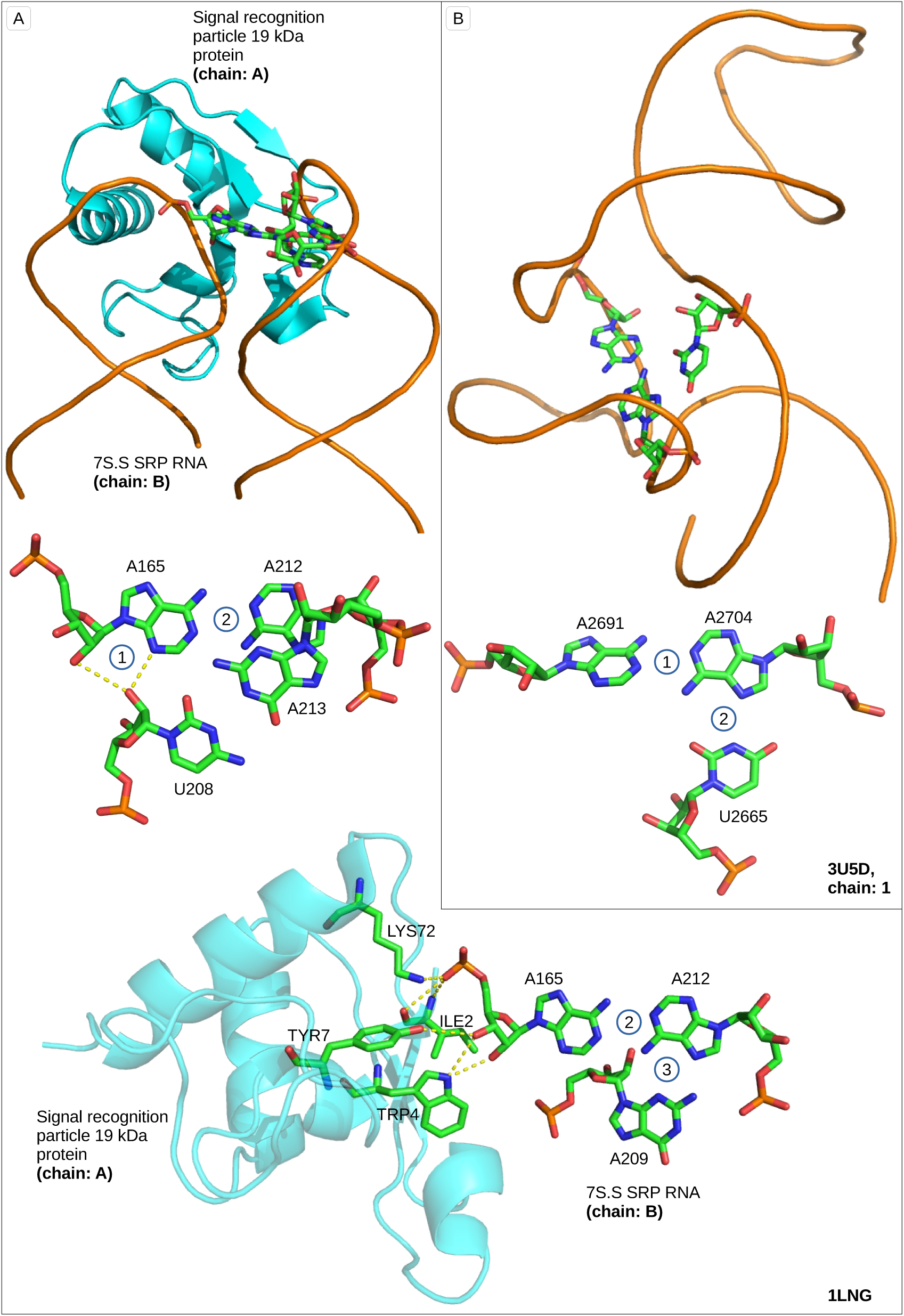
Two examples of base triples containing A:A W:W Trans pair have been shown — **(A)** a kissing loop interaction observed in SRP RNA S domain of *M. jannaschii* and **(B)** a junction loop motif observed in 25S rRNA of *S. cerevisiae.* In the first example, different intermolecular interactions have been annotated as, (1) Type II A-minor interaction (2) A212(B):A165(B) W:W Trans and (3) A212(B):G209(B) H:S Trans. In the second example, different intermolecular interactions have been annotated as, (1) A2691(1):A2704(1) W:W Trans and (2) U2665(1):A2704(1) W:H Trans

#### 3.4.1 Base triples involving A:A W:W Trans: energetics and context analysis

Apart from being a component of base quartets, base triples occur independently in the structurally conserved homologous positions in 23S rRNA, 25S rRNA, Group I intron and SRP 7S.S RNA as a part of different types of loops, such as kissing loop (context 1,4 and 11 in Table S8), junction loop (context 1, 5 and 9 in Table S8 and context 1, 2 of Table S9) or internal loop (context 2, Table S9) motifs. These structural domains have further biophysical roles as elaborated in Figure 5. We have found that, in SRP19-7S.S SRP RNA complex of *Homo sapiens* (eukaryote), *M. jannaschii(archaean)* and *Sulfolobus solfataricus* (archaean), the SRP19 protein binds to a kissing loop region of the RNA (Figure 5 A), which is composed of a conserved A:A-G triple (context 11 of Table S8). This A:A-G triple involves a A:A W:W Trans base pair. Interestingly, one of the adenines of this base pair further participates in a type II A-minor interaction and also interacts with the SRP19 protein from its sugar-phosphate backbone (Figure 5 A).

Figure 6 shows the ground state optimized geometries and intrinsic interaction energies for the frequently occurring base triples, involving A:A W:W Trans base pairs. Please note that, when a third base (U and G respectively) pairs with the Hoogsteen edge of the central adenine, the hydrogen bonding interaction mediated through N6 gets improved, whereas the hydrogen bonding interaction involving N1 gets compromised. This is evident from the change in hydrogen bond donor-acceptor distances as shown in Figure 6 and change in the red shift values of the N-H bond stretching frequencies reported in Table 1. Pairing of the Watson-Crick edge of uracil with the Hoogsteen edge of the central adenine in A:A-U triple results in a planar triple (Figure 6A, Table 2), which has a significantly higher interaction energy compared to the nonplanar A:A-G triple (Figure 6B, Table 2). In the A:A-G triple, sugar edge of guanine pairs with the Hoogsteen edge of the central adenine. It is to be noted that, interaction energy of isolated A:A-U (-21.9 kcal/mol) and A:A-G (-19.5 kcal/mol) triples are even lower than isolated G:G W:W Trans (-25.8 kcal/mol) or canonical G:C base pairs. This indicates high overall flexibility of the structural motifs which include these relatively easily breakable triples. Although, the A:A-U (Figure 6A) and A:A-G (Figure 6B) triples occur independently, the A:A-C triple (Figure 6C and D) occur only as a part of a A:A-C-U base quartet (Figure 7A). In the A:A-C triple, the Watson-Crick edge of cytosine pairs with the Hoogsteen edge of the central adenine in *trans* orientation. As discussed in our earlier works,^67,68^ such a geometry of interaction between WC edge of cytosine and Hoogsteen edge of adenine is possible only if either of cytosine N3 and adenine N7 gets protonated. We have considered both the possibilities and have observed that (Figure 6 C, D, Table 2), N3 protonated cytosine results in a nonplanar triple with lower interaction energy (-32.1 kcal/mol) whereas, the triple with adenine N7 protonation (-49.3 kcal/mol) is more stable. This is due to the fact that, although cytosine N3 is easier to protonate than adenine N7 protonation in solution,^68^ protonation induced charge redistribution in cytosine disfavors hydrogen bonding interactions via O2 of cytosine. On the other hand, protonation at N7 of adenine favors the hydrogen bonding interactions via N6 of adenine^68^. Higher charge dipole interaction between neutral cytosine and protonated adenine, compared to the same between charged cytosine and neutral adenine provides another explanation for such observations^67^. Therefore, in the next section, we have considered only the N7 protonation of adenine while discussing the A:A-C-U quartet.

**Figure 6.**
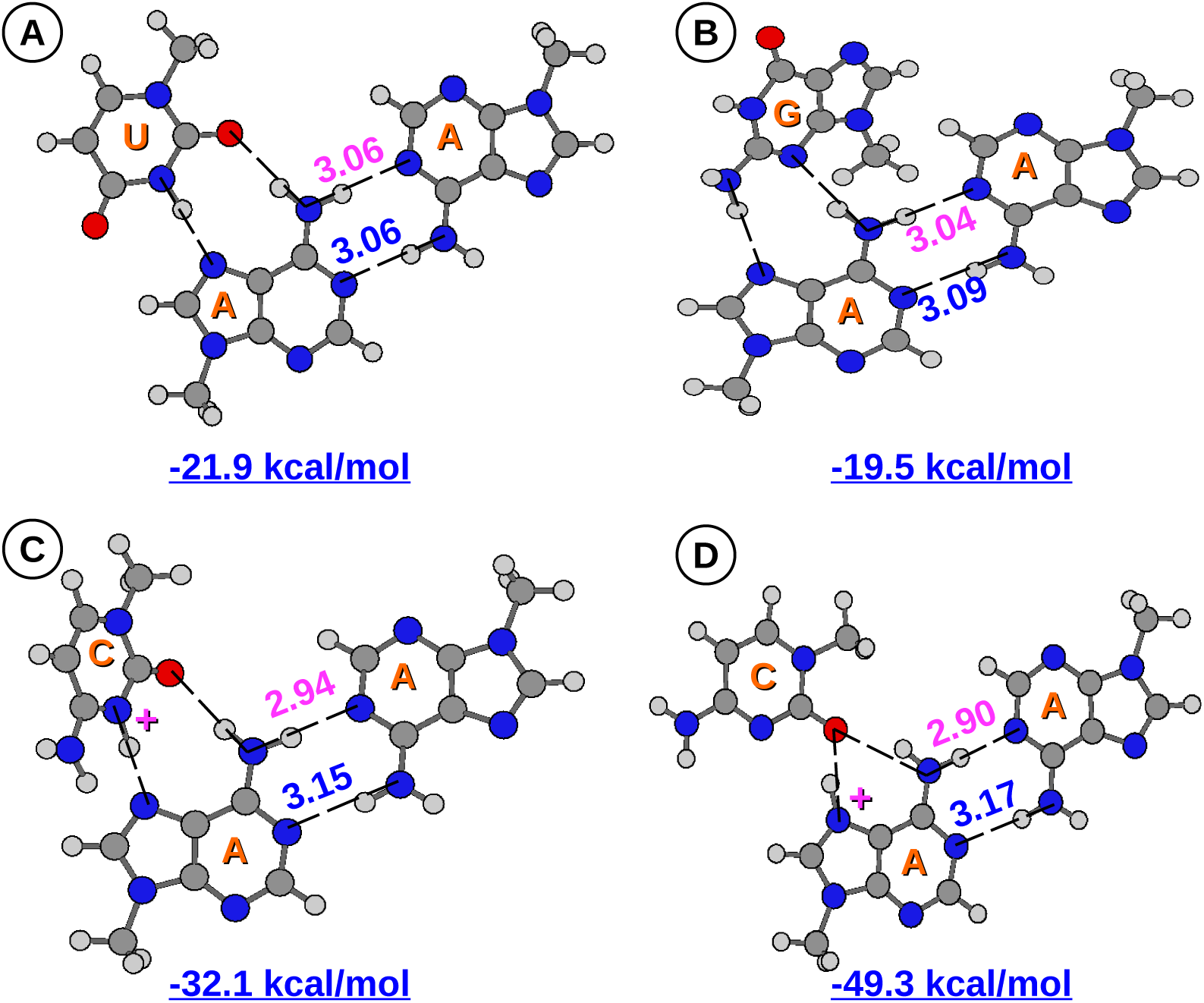
Ground state optimized geometries and interaction energies at M05-2X/6-31G++(2d,2p) level of theory have been shown for **(A)** A:A-U, **(B)** A:A-G and **(C,D)** A:A-C triples. A:A-C triple has been modeled in two ways **(C)** with N3 protonated cytosine and **(D)** with N7 protonated adenine. Hydrogen bond donor-acceptor distances (in Å) are shown for the two N-H…N type hydrogen bonds of the A:A W:W Trans pair.

**Figure 7.**
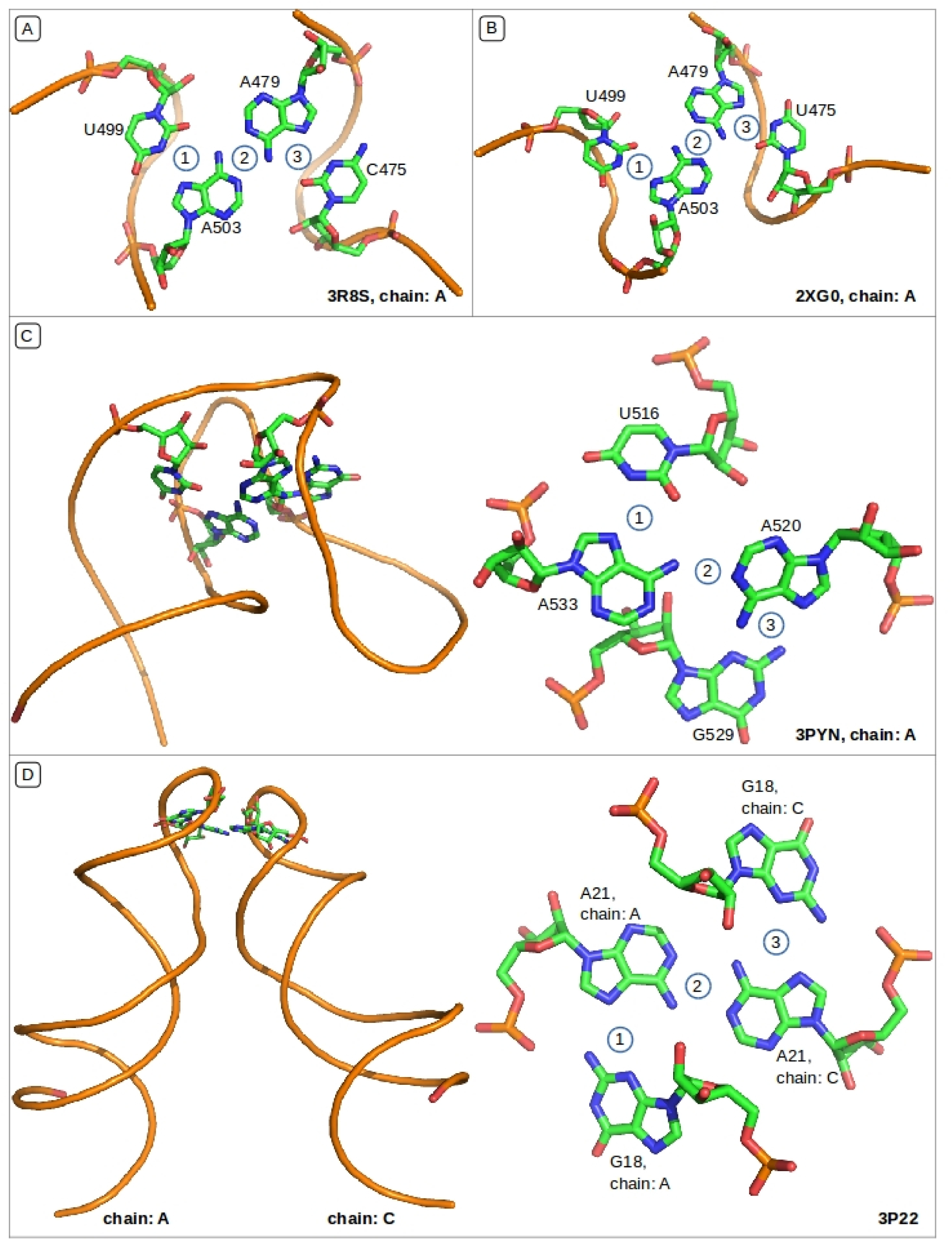
**(A)** 23S rRNA *E. coli,* loop-loop interaction 1) U499(A):A503(A) W:H Trans 2) A503(A):A479(A) W:W Trans 3) A479(A):C475(A) W:H Trans; **(B)** 23S rRNA *T. thermophilus,* loop-loop interaction 1) U499(A):A503(A) W:H Trans 2) A503(A):A479(A) W:W Trans 3) U479(A):A475(A) W:H Trans; **(C)** 16S rRNA *T. thermophilus,* internal loop 1) U516(A):A533(A) W:H Trans 2) A533(A):A520(A) W:W Trans 3) A520(A):G529(A) H:S Trans; **(D)** Synthetic core ENE hairpin from KSHV PAN RNA, kissing loop interaction 1) A21(A):G18(A) H:S Trans 2) A21(A):A21(C) W:W Trans 3) A21(A):G18(C) H:S Trans;

**Table 2.** Inter base pair rotational parameters (Buckle, Open and Propeller) and translational parameters (Stagger, Shear and Stretch), as calculated by the NUPARM package, have been reported for the base pairs present in the optimized geometries of base triples and quartets containing A:A W:W Trans base pair.

| System | Base Pair | Buckle | Open | Propeller | Stagger | Shear | Stretch |
| --- | --- | --- | --- | --- | --- | --- | --- |
| **Triples** |  |  |  |  |  |  |  |
| A:A-G | A:A W:W Trans | 7.29 | 0.94 | 8.56 | -0.15 | 2.4 | 3.05 |
|  | A:G H:S Trans | 8.28 | 75.29 | 45.41 | 0.74 | 0.03 | 3.73 |
| A:A-U | A:A W:W Trans | -0.01 | 0.07 | -0.03 | 0 | 2.28 | 3.06 |
|  | U:A W:H Trans | 0.04 | 4.93 | 0 | 0 | 0.07 | 2.81 |
| A:A(+)-C | A:A W:W Trans | 6.65 | 5.35 | -10.95 | -0.18 | 2.23 | 3.04 |
|  | C:A W:H Trans | 25.26 | 2.99 | 53.88 | 0.47 | 0.51 | 2.61 |
| A:A-C(+) | A:A W:W Trans | 1.3 | 6.8 | 4.67 | -0.02 | 2.26 | 3.03 |
|  | C:A W:H Trans | -2.14 | -29.81 | -2.04 | -0.02 | -1.75 | 3.07 |
| **Quartets** |  |  |  |  |  |  |  |
| A:A-G-G | A:A W:W Trans | 51.94 | 0.01 | 23.16 | 0 | 2.63 | 3.02 |
|  | A:G H:S Trans | 9.65 | 74.49 | 47.91 | 0.83 | 0.02 | 3.72 |
|  | A:G H:S Trans | 9.62 | 74.57 | 47.91 | 0.83 | 0.02 | 3.72 |
| A:A-G-U | A:A W:W Trans | 9.47 | 1.44 | 7.9 | -0.12 | 2.36 | 3.07 |
|  | A:G H:S Trans | 8.07 | 75.07 | 45.82 | 0.74 | 0.03 | 3.73 |
|  | U:A W:H Trans | 0.83 | 4.94 | -1.64 | -0.01 | 0.08 | 2.81 |
| A:A-C(+)-U | A:A W:W Trans | 3.98 | 6.13 | -7.21 | -0.21 | 2.2 | 3.05 |
|  | C:A W:H Trans | 30.35 | 2.11 | 56.95 | 0.47 | 0.58 | 2.6 |
|  | U:A W:H Trans | -2.71 | 4.62 | -0.18 | -0.01 | 0.12 | 2.82 |
| A:A(+)-C-U | A:A W:W Trans | 2.09 | 7.63 | 4.04 | -0.01 | 2.22 | 3.04 |
|  | C:A W:H Trans | -4.14 | -30.68 | -4.66 | -0.06 | -1.74 | 3.1 |
|  | U:A W:H Trans | 0.46 | 4.55 | -0.18 | 0.01 | 0.13 | 2.82 |

#### 3.4.2 Base quartets involving A:A W:W Trans: energetics and context analysis

Structural alignment of homologus RNAs from different species shows that, the base quartets involving A:A W:W Trans remain conserved across species (Table S8). However, an exception is observed in 23S rRNA of *H. marismortui* and *E. coli* (context 8 of Table S8), where the uracil of a A:A-U-U quartet co-varies with cytosine. As shown in Figure 7A and B, this quartet mediates a tri-loop tri-loop interaction and therefore, has significant structural roles^69,70^. Interestingly, such substitution of uracil by cytosine involves protonation of either adenine N7 or cytosine N3, which are thermodynamically unfavorable processes under physiological pH ( ~7.4)^71^. However, the energetic cost perhaps gets compensated by the high interaction energy (-63.0 kcal/mol) of the resulting A:A-C-U quartet (Figure 8B). The other two quartets, A:A-G-U (Figure 8C) and A:A-G-G (Figure 8D), have lower interaction energies and are comparatively less planar than the A:A-U-U and A:A-C-U quartets, as denoted by high Buckle, Open and Propeller values of the corresponding constituent A:G H:S Trans base pairs (Table 2). It is important to note that, except for only two instances, formation of base quartet significantly increases the hydrogen bond donor-acceptor distances of the central A:A W:W Trans base pair compared to isolated A:A W:W Trans pair where the hydrogen bond donor-acceptor distance is 3.5 *Å* (Figure 8). We also observe a decrease in the red shift values of the symmetric stretching frequency of the corresponding hydrogen bond donors (Table 1). Therefore, the N-H…N hydrogen bonds of the central A:A W:W Trans pair become weaker and more flexible within the quartets. Participation of such fragile base quartets in connecting distant structural motifs (Figure 7) are conserved across species, e.g., context 1, 3 and 6 for A:A-G-U quartets (Table S8), context 1, 7 and 10 for A:A-G-G quartets (Table S8), etc. The indicated overall flexibility of domains containing these ‘fragile’ quartets, may be implicated in the functional dynamics, and have been discussed later in the context of NMR structures.

**Figure 8.**
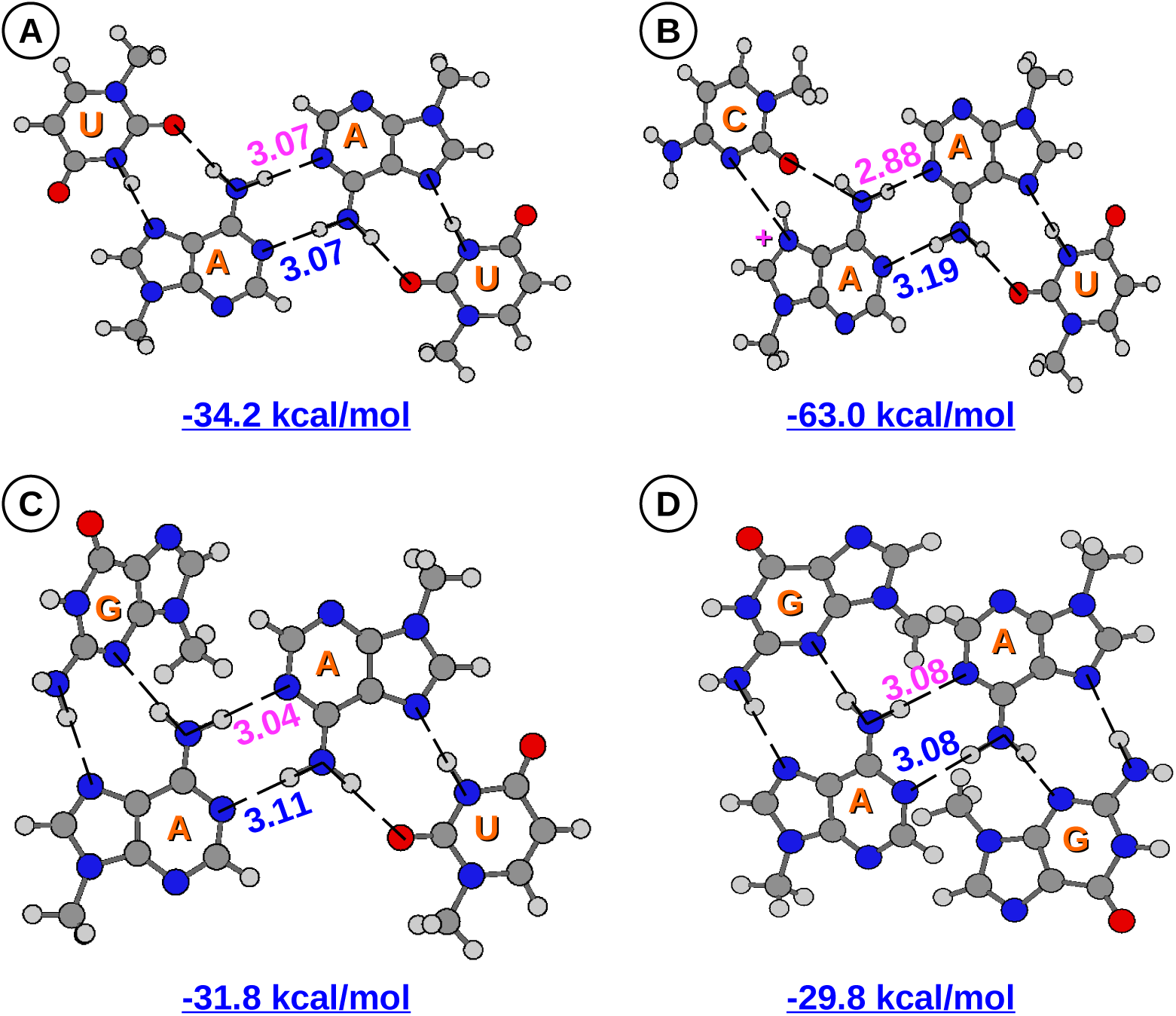
Ground state optimized geometries and interaction energies at M05-2X/6-31G++(2d,2p) level of theory have been shown for **(A)** AAUU, **(B)** AACU, **(C)** AAGU and **(D)** AAGG quartets. AACU triple has been modeled with N7 protonated adenine forming A(+):C H:W Trans base pair. Hydrogen bond donor-acceptor distances (in Å) are shown for the two N-H…N type hydrogen bonds of the A:A W:W Trans pair.

### 3.5 Why G:G W:W Trans pairs cannot form base-triples/quartets like A:A W:W Trans pairs?

The general observation emerging from the above discussions is that, purine-purine W:W Trans base pairs occur in backbone stretches characterized by low *ρ_min_* values (Figure 2b), when they are part of a Sharp-turn, and are not involved in any higher order interactions. On the other hand, they populate backbone stretches characterized by high *ρ_min_* values (Figure 2b), when they are involved in higher order interactions and participate in formation of larger structural motifs. Thus, absence of G:G W:W Trans pairs in the latter category (Table S6) indicates that they can not engage in higher order interactions such as triples and quartets. This observation explanation can also be substantiated on the basis of QM results.

We have observed only three types of base triples which involve A:A W:W Trans pair, A:A-U, A:A-C and A:A-G. However, the equivalent G:G-U, G:G-C or G:G-G triples having G:G W:W Trans pairs in the place of A:A W Trans, are absent. Absence of G:G-U and G:G-C triples can be explained from the fact that, free hoogsteen edges of G:G W:W Trans lacks the complementary hydrogen bonding network with respect to Watson Crick edge of uracil and protonated cytosine. Therefore U:G W:H Trans and C(+):G W:H base pairs equivalent to U:A W:H Trans (as present in A:A-U) and C(+):A W:H Trans (as present in A:A-C) pairs, respectively, can not be formed. However, the A:A-G triple is composed of a G:A S:H Trans base pair and an equivalent G:G S:H Trans base pair is possible due to presence of complementary hydrogen bond donor acceptor network between the interacting edges.^5^ Non occurrence of the G:G-G triple therefore requires a different explanation. Hence, we have calculated the ground state electronic properties of a modeled G:G-G triple to have a better insight. We modeled the geometry of putative G:G-G triple by superposing first guanines of G:G W:W Trans and G:G H:S Trans base pairs as shown in Figure S6 and found that the modeled geometry of G:G-G triple converges to a stable minima with high interaction energy, −37.7 kcal/mol (Figure S6). Again, we have found that G:G H:S Trans base pair is inherently unstable since, on full geometry optimization, the G:G H:S Trans base pair converges to a geometry which is remarkably different from its native crystal geometry (Figure S5). Clearly the rationalization of the observation does go beyond simple considerations of energetics. One possibility may be the evolutionary pressure related to functional requirements. Our QM calculations suggest that, the N-H…O type hydrogen bonds of G:G W:W Trans are significantly stronger than N-H…N type hydrogen bonds present in A:A W:W Trans pair. Red shift of the symmetric stretching of amino group's N-H bonds is 215.5 cm^−1^ higher for N-H…O bonds of G:G W:W Trans (Table 1). With these strong hydrogen bonds, G:G W:W Trans pair is inherently rigid by design as compared to A:A W:W Trans pair. The modeled G:G-G triple also has a high interaction energy compared to natural A:A W:W Trans containing triples (Figure 6). Therefore, structural motifs which require flexibility in its design, do not contain G:G W:W Trans base pairs. As discussed earlier, RNA structural motifs where the A:A W:W Trans containing base triples and quartets are conserved across species, appear to be flexible by design and therefore do not possess G:G W:W Trans base pairs.

### 3.6 Possible A:A W:W Trans mediated conformational dynamics in RNA

Structural alignment studies of 23S rRNA crystal structures reveal a conserved internal loop consisting of A:A base pair interacting with minor grove side of G:C helix. Out of 9 crystal structures of 23S rRNA in the non-redundant dataset, 6 have A:A base pair in W:W Trans geometry and 3 have A:A base pair in W:H Trans geometry (Table S6). Interestingly, in the latter case, the second adenine participates in type I A-minor interaction with G:C helix (Figure 9 C and D). A-minor interactions are considered to be the most abundant and functionally significant long-range interactions stabilizing tertiary structure in large RNAs^28^.

**Figure 9.**
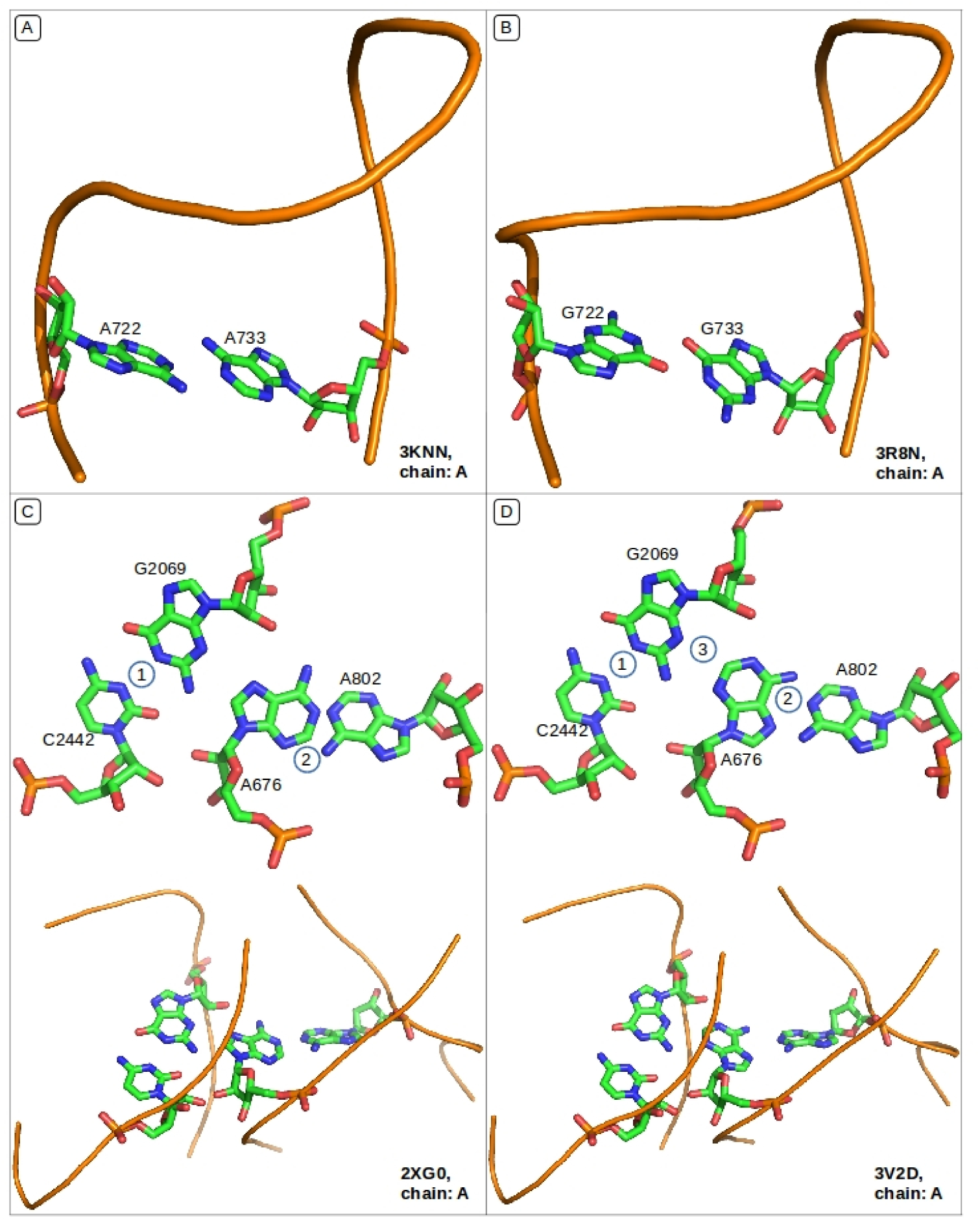
**(A)** 16S rRNA *T. thermophilus*, A722(A):A733(A) W:W Trans; **(B)** 16S rRNA *E. coli*, G722(A):G733(A) W:W Trans; **(C)** 23S rRNA *T. thermophilus,* 1) G2069(A):C2442(A) W:W Cis 2) A676(A):A802(A) W:W Trans; **(D)** 23S rRNA *T. thermophilus*, 1) G2069(A):C2442(A) W:W Cis 2) A676(A):A802(A) W:H Trans 3) Type I A-minor interaction;

The above observation is indicative of possible dynamics at play involving ~ 180 degree rotation about the glycosidic bond leading to transition of A:A base pair geometry from W:W Trans to W:H Trans and vice-versa. This rotation also takes care of the requirement of the antiparallel local strand orientation. It is to be noted that, in both the W:W Trans and W:H Trans geometries the A:A base pair maintains an anti-parallel local strand orientation, which is possible since one of the bases is in *syn* conformation in W:W Trans geometry. Importance of this evidence is further highlighted by the fact that *syn* nucleobases have earlier been shown to occur widely in regions of functional importance in a variety of RNAs^72^. One should not expect similar W:W Trans to W:H Trans switching via rotation about the glycosidic bond in the case of G:G base pairs, since the interaction energy difference between G:G W:W Trans and G:G W:H Trans base pairs are significantly higher compared to that for A:A W:W Trans and A:A W:H Trans base pairs^31^.

### 3.7 Conditional covariation of A:A W:W Trans with G:G W:W Trans

Base pairs with similar *C*1′ — *C*1′ distance and same glycosidic bond orientation are expected to be isosteric, and hence covary, without significantly disturbing the existing local interactions. Thus, in principle, they can replace each other without causing any susbtantial change to local three-dimensional geometric paths and sugar-phosphate backbone orientations^16,59^. However, A:A and G:G W:W Trans pairs do not generally covary with each other. One instance where we have noticed A:A W:W Trans being replaced by G:G W:W Trans (Figure 9) is in 16S rRNA of *T. thermophylus* (A722:A733, 3KNN, chain A) and E. *coli* (G722:G733, 3R8N, chain A) where, both the base pairs are a part of a bulge loop and have anti-parallel local strand orientation since one of the bases is in *syn* conformation. In neither of these instances do we observe the purine-purine base pairs to be invoking their respective characteristic, and remarkably different, physicochemical surfaces (see Figure S9) to interact with ions, proteins or participate in higher order tertiary interactions. Other instances of A:A and G:G W:W Trans pairs are respectively associated with their own characteristic functionally important interactions with nearby nucleotides and amino acids, even when they occur as a part of Sharp-turn and hence do not covary.

### 3.8 Role of A:G W:W Trans type recurrent hydrogen bonded interactions in functional RNAs

Among the 6 A:G W:W Trans type recurrent hydrogen bonded interactions, which have been detected using BPFIND software (Table S11 and Figure S7), 4 are found in 16S rRNA, one is found in 25S rRNA and other is found in 23S rRNA. In 16S rRNA, the A:G W:W Trans pair is detected between A1492 and G530. Interestingly, although it has been known since long^73^ that ribosome differentiates between the cognate and near cognate tRNA binding with the help of these two residues (along with A1493),^6^ the A:G W:W Trans geometry has not formally been recognized as a base pair. Here we have attempted to explain how characteristic geometry and stability of A:G W:W Trans type recurrent hydrogen bonded interactions facilitate its occurrence in this specific functional context. At the same time, we are also able to figure out the role of these base pairs in other structural and functional contexts, such as, mediating loop - loop interactions in in 25S rRNA and forming bifurcated triple in 23S rRNA.

#### A:G W:W Trans in 16S rRNA

As shown in Figure 10 B, in the bound state, A1492 gets engaged in type II A-minor interaction^28^ with mRNA codon base and G530 gets engaged in type II G-minor interaction^77^ with tRNA anticodon base. N3 and O2^'^ atoms of both the purines act as ‘clips’ for the O2′ atoms of canonically paired codon(mRNA):anticodon(tRNA) bases. These interactions are shown in yellow broken lines in Figure 10 B. Here, A:G W:W Trans pair which is intrinsically non-planar, is in a conformation which is tailor-made for discrimination against near isosteric base pairs like G:U W:W Cis and other noncanonical base pairs^78^. This unique geometry is characterized by large *C*1′-*C*1′ distance (~12.9 Å), buckle (~24°) and propeller twist (~40°). The formation of A1492:G530 W:W Trans pair in cognate tRNA bound 16S rRNAs is further supported by the interactions of A1492 with C518 and Ser50 of protein S12.

**Figure 10.**
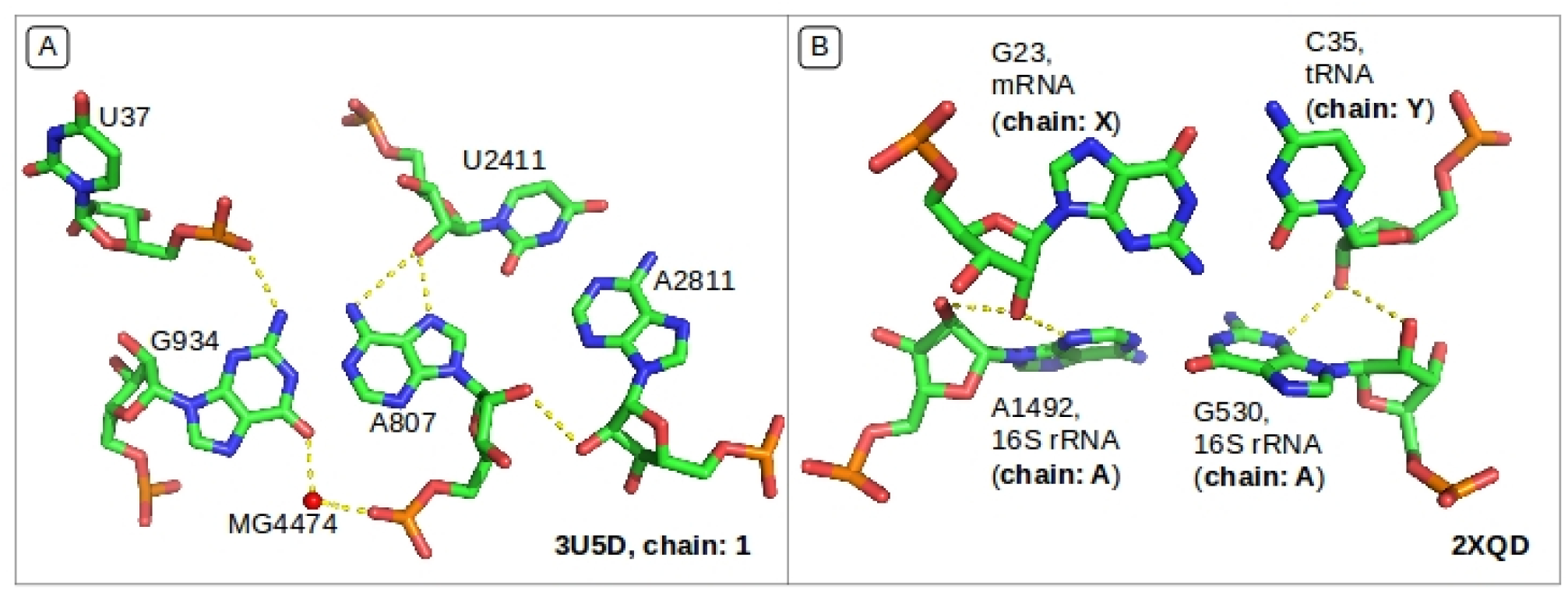
**(A)** A807:G934 W:W Trans, part of internal loop interacts with canonical A2811:U2411 W:W Cis (helix) and U37 of an hairpin loop; **(B)** 1492A:530G W:W Trans interacting with second codon:anticodon base pair via type II A-minor and G-minor interactions respectively.

#### A:G W:W Trans in 25S rRNA

Figure 10 A shows the detected occurrence of A:G W:W Trans in 25S rRNA of *S. cerevisiae.* This base pair is a part of an internal loop, where the guanine adopts a *syn* conformation, and is therefore able to avoid any sharp turn in the backbone, by maintaining the required anti-parallel local strand orientation. Interestingly, this particular A:G W:W Trans base pair mediates interaction of the internal loop with another hairpin loop and a canonical helix located in the vicinity. Interaction of the A:G pair with the minor groove of helix (via A2811:U2411 canonical pair) and U37 of the hairpin loop is an integral part of the design of the tertiary contact (Figure 10 A), which is possibly uniquely provided by the A:G W:W Trans base pair.

#### A:G W:W Trans in 23S rRNA

The one instance observed in 23S rRNA of *T. thermophilus* (PDB Id: 3UZ8) is a part of a bifurcated triple (example 6 of Figure S8). Adenine being the central base of the bifurcated triple, the A2267:G2271 W:W Trans pair is highly deviated from the planarity, as suggested by the corresponding high E-value (1.32). The high RMSD value between the corresponding F_*opt*_ and H_*opt*_ geometries (0.81 JÅ) is also suggestive of an important role of the bifurcated triple environment in the significant distortion of the H_*opt*_ structure compared to the isolated F_*opt*_ optimized structure.

### 3.9 Purine-Purine W:W Trans pairs in solution NMR structures

In the 591 solution NMR structures studied in this work, we have identified 17 instances of A:A W:W trans and 9 instances of G:G W:W Trans. However, no instances of A:G W:W Trans has been detected. Structures containing these base pairs include GAAA tetraloop receptor dimer, HIV dimerization initiation site, HIV-1 Rev binding RNA, etc.

In the GAAA tetraloop receptor dimers (Table S19), the A:A W:W Trans pair is a part of A:A-U triple that mediates binding of the hairpin GAAA loop of one monomer with the receptor internal loop of the other (Figure 11F)^79–81^. In HIV dimerization initiation site (Context 1 of Table S20) the dimerization of two monomers takes place via a kissing loop interaction^82^. Structural alignment of different models reported in PDB 2D19 suggest that, the base pairing interaction between A13 and A15 (an fundamental component of the kissing loop motif) fluctuates between W:W Cis and W:W Trans geometries. As shown in Figure 11E, A13:A15 W:W Trans pairing results in a Sharp-turn motif (Model 1 in 11E) which disappears when the geometry switches to A13:A15 W:W Cis (Model 2 in 11E). This is supportive of the inherent flexible nature of the A:A W:W Trans base pair as suggested by our QM calculations.

**Figure 11.**
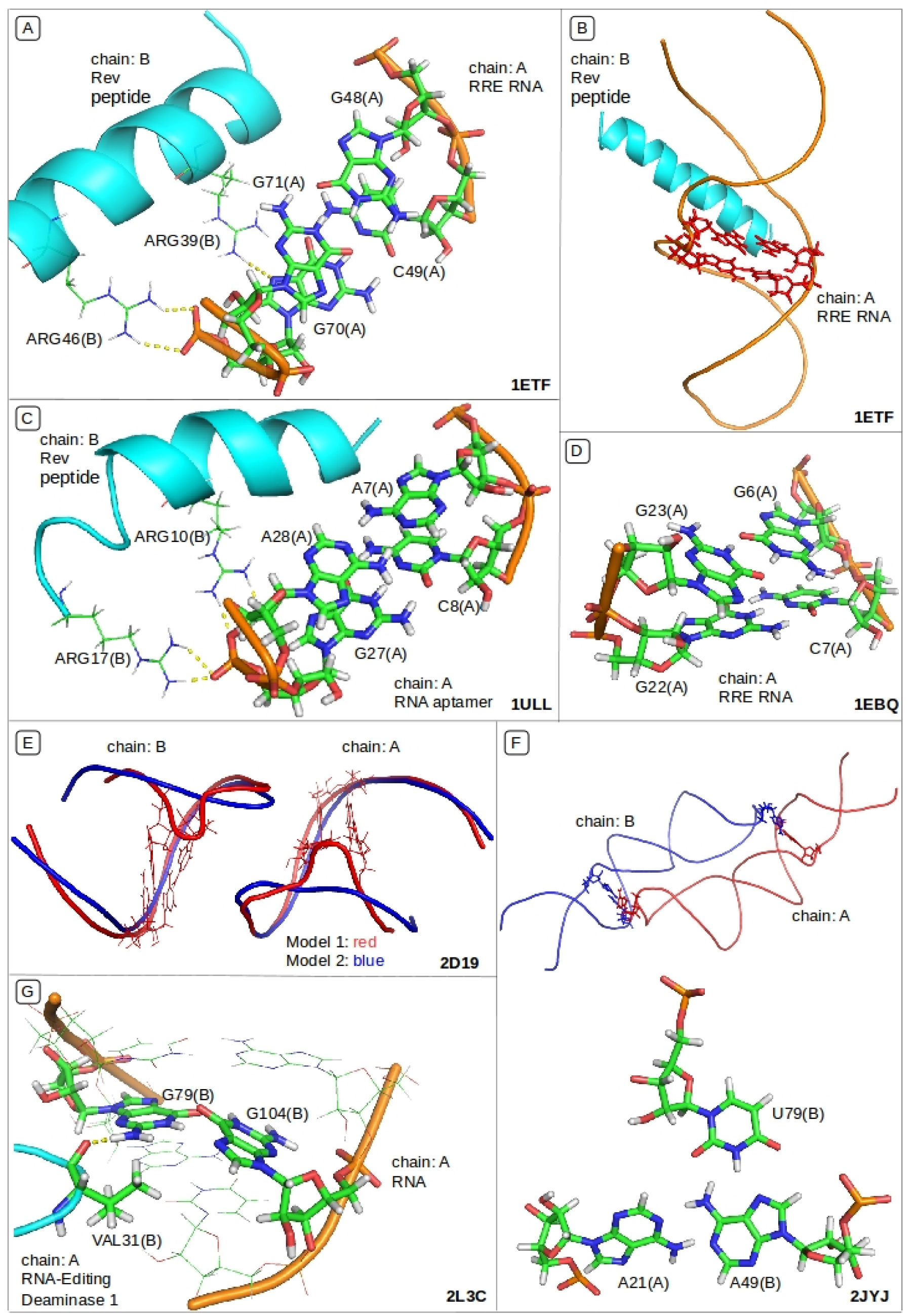
Structural contexts of purine-purine W:W Trans pairs detected in solution NMR structures of RNA. (A & B) HIV-1 Rev peptide-RNA aptamer complex with Sharp-turn composed of G:G W:W Trans at the bonding site (1ETF), (C) the same with Sharp-turn composed of A:A W:W Trans (1ULL), (D) free RRE-IIB with no Sharp-turn (1EBQ), (E) Kissing loop at HIV dimerization initiation site (2D19) with (red) and without (blue) Sharp-turn, (F) GAAA tetraloop receptor dimer (2JYJ) with A:A-U triple mediated loop-loop interaction and (G) G:G W:W Trans pair within a canonical helix (2L3C)

HIV-1 Rev peptide-RNA aptamer complex exhibits a deep penetration of an alpha-helix into a widened RNA major groove (Figure 11B)^27,83,84^. Analysis of the NMR structures reveal that, the distinctive binding pocket for the peptides in stem IIB of the Rev responsive element (RRE-IIB) of HIV-1 mRNA is composed of a Sharp-turn motif consisting of adjacent G:C W:W Cis and G:G W:W Trans pairs (Context 2 of Table S20). Figure 11A shows composition of the Sharp-turn and interaction of corresponding backbone atoms with the Rev peptide. Comparison of the free (1EBQ, 1EBR) and bound (1EBS) RNA structures also reveal that, in the free RNA, one of the guanines is present in *syn* conformation and therefore the resulting G:G W:W Trans pair does not produce a Sharp-turn in the backbone (Figure 11D)^83^. Interestingly, in another structure of HIV-1 Rev peptide-RNA aptamer complex (1ULL), which is in-vitro selected for high affinity binding, the Sharp-turn is composed of A:A W:W Trans pair (Figure 11C)^27^. It is noteworthy that, as shown in Figure 11A, only the backbone atoms of the Sharp-turn are involved in further nonbonding interactions and therefore, a covariation between isosteric A:A and G:G W:W Trans pairs are allowed.

G:G W:W Trans base pair with one guanine in *syn* conformation (G104(B) in Figure 11G) is also observed in the ADAR2 dsRBM-RNA complex, where the G:G pair fits within an otherwise canonical helix (Table S20)^85^. Another example of G:G W:W Trans is found in influenza A, which is an RNA virus with a genome of eight negative sense segments (Context 5 of Table S20). Segment 7 mRNA contains a 3′ splice site for alternative splicing to encode the essential M2 protein. In the chain B of PDB 2L3C, G32:G10 W:W Trans along with A9:U33 W:W Cis constitute a Sharp-turn motif which is located next to the splicing site (between G10 and G11 in the internal loop). Interestingly, this Sharp-turn between G10 and A9 is conformationally dynamic and is absent in Model 5 and Model 12 (PDB: 2L3C) where the sharp turning of the backbone has been avoided as the G10 residue is flagged out in Model 5 and G10 is present in *syn* conformation in Model 12^86^.

## 4 Summary and Conclusion

In this work, we have used a combination of bioinformatics and QM methods to investigate the physicochemical features of purine-purine W:W Trans base pairs. With the largest C1′-C1′ distance and, usually, a parallel local strand orientation, purine-purine W:W Trans base pairs constitute an important class of base pairs in RNA. To accommodate such base pairs in an otherwise double helical environment, either the RNA backbone has to adopt a special fold or one of the bases has to be present in *syn* orientation. Prevalence of such *syn* nucleobases in the active sites of functional RNAs has been reported by Bevilacqua and co-workers^72^.

Parameters characterizing the local backbone geometry of purine purine base pairs, observed in our dataset, indicated a bimodal distribution. The first mode is characterized by occurrence of a A:A/G:G W:W Trans pair adjacent to a canonical A:U/G:C W:W Cis pair, resulting in a sharp turn in one of the two strands. We have identified such structural contexts as ‘Sharp-turn’ motifs and have studied their conservation and covariation patterns. Structural alignment of homologous locations of different RNA chains suggests that such Sharp-turn motifs constitute distinct sets corresponding to those with A:A or its variants, and G:G or its variants, respectively, and are conserved across species. Notably, within each set there are some compositional variations observed, primarily due to (a) the substitution of one or both of the purines of W:W Trans pair with pyrimidines, or, (b) covariation between A:U and G:C canonical pairs at the adjacent position. Such compositional variations notwithstanding, for a given context, the non-bonding interactions associated with these Sharp-turns remain conserved. Further, though otherwise isosteric, A:A and G:G W:W Trans base pairs do not generally covary within any given context. This can be easily explained based on the differences in their respective physicochemical surfaces, and the non-bonding interaction network they may participate in. We also have reported association of these Sharp-turns with binding of ribosomal proteins and ligands and have suggested the possibility of the co-evolution of ribosomal protein S16 and 16S rRNA.

In the second mode, no such sharp turning of the backbone is observed and it is specific to only A:A W:W Trans pairs. Interestingly, all the occurrences of A:A W:W Trans base pairs in this mode are also part of base quartets or independent base triples, where the WC edge of uracil/cytosine or the sugar edge of guanine interact with the free Hoogsteen edges of A:A W:W Trans in *trans* orientation. Our context analysis of such triples/quartets reveals that, they mediate between distant structural folds via loop-loop interactions, which are conserved across species. In some cases such structural folds provide the binding pocket for proteins. We have found that, G:G W:W Trans pairs do not form base triples via their free Hoogsteen edges with uracil/cytosine, due to lack of complementary hydrogen bond donor-acceptor network. However, given the fact that, both the G:G H:S Trans (although intrinsically unstable) and G:G W:W Trans base pairs occur in RNA crystal structures, we have modeled a G:G-G triple by superposing the first guanines of both the G:G H:S Trans and W:W Trans base pairs, respectively. This modeled G:G-G triple is intrinsically stable but is still not observed in nature. This apparent discrepancy may be rationalized by considering the fact that, inter-base hydrogen bonds in A:A W:W Trans pair gain greater flexibility on triple/quartet formation and thus are possibly designed to provide an overall flexibility to the larger motif. In the context of flexible motifs, possibly designed for special functional requirements, this intrinsic difference in structural features may be suggestive of evolutionary pressure in favor of the more flexible A:A W:W Trans pair over the relatively rigid G:G W:W Trans pair. Similar trends are also observed in the available solution NMR structures of RNA.

We have also identified occurrences of recurrent hydrogen bonded interactions between A1492 and G530 in W:W Trans geometry in 16S rRNA bound to the cognate tRNA at A-site. Comparison of the H_*opt*_ and F_*opt*_ geometries of these instances confirm that these occurrences are not fortuitous. It must be noted that, participation of these A1492 and G530 residues along with A1493 residue in discriminating canonical codon:anticodon pairs from nearly isosteric pairs like G:U W:W Cis has already been discussed in literature^73,76,78^. In this work, we underline the fact that, the unique geometry corresponding to the interaction between A1492 and G530 in W:W Trans geometry (characterized by the C1′-C1Å distance of ~12.9 Å and average Buckle and Propeller of ~24° and ~40°, respectively) enables A1492 and G530 to discriminate the canonical codon:anticodon pairs from the others. We are also able to identify structural and functional roles of A:G W:W Trans pairs in 23S and 25S rRNAs.

To sum up, in this work we have systematically highlighted the versatile roles of purine-purine reverse Watson-Crick base pairs in the context of RNA structure and dynamics. We have demonstrated that, (i) occurrences of A:A/G:G W:W Trans pairs adjacent to a canonical pair causes a sharp turn in one of the backbone strands, (ii) the Sharp-turn motif has several biophysical roles including formation of the binding pocket for Rev peptides in stem IIB of the Rev responsive element of HIV-1 mRNA, (iii) in the Nature, the intrinsic flexibility of A:A W:W Trans is possibly implemented in the design of larger motifs with overall flexibility, (iii) A:A W:W Trans and W:H Trans geometries exist in a dynamic equilibrium and (iv) unique buckled and twisted geometry of A:GW:W Trans pair is necessary for the recognition of cognate tRNA by the 30S ribosomal subunit. These information will on one hand help to bridge the gap between the sequence, structure and function domains of RNA and on the other, will aid the bio-engineering community in designing oligonucleotides with desired structure and function.

## 5 Acknowledgement

AH and SB are thankful to CSIR and UGC, respectively for Senior Research Fellowship. DB thanks partial financial support from BARD project of Department of Atomic Energy, Government of India. AM thanks DBT, Government of India project BT/PR-14715/PBD/16/903/2010 for partial funding and financial support.

## Footnotes

1 In this work, to annotate the base pairing interactions we have followed a nomenclature which is slightly different from the Leontis-Westhof^15^ nomenclature. If the edge X of base A interacts with the edge Y of base B in cis (or *trans)* orientation, that is annotated as A:B X:Y Cis (or, A:B X:Y Trans).

2 Note that, due to lack to planarity and presence of only one strong hydrogen bond, it is arguable whether the recurrent hydrogen bonding interactions observed between adenine and guanine in W:W Trans orientation are to be considered as base pair or not. However, in this work, for the sake of generality we will refer to such recurrent interactions as A:G W:W Trans.

3 *C*1′ − *C*1′ distances have been calculated for the optimized geometry modeled with both the sugars in *anti* conformation

4 We have calculated the average values (13.5 Å for A:A, 13.4 Å for G:G and 12.9 Å for A:G) and corresponding standard deviations (0.3 Å for A:A, 0.3 Å for G:G and 0.2 Å for A:G) of the C1′-C1′ distance for all the instances of A:A, G:G and A:G W:W Trans base pairs detected in the complete dataset.

5 11 instances of G:G H:S Trans in HDRNAS dataset have been reported in RNABP COGEST.

6 A1492 and G530 (along with A1493) are conformationally active, universally conserved, essential nucleotides of 16S rRNA in ribosome crystal structures that are bound with cognate transfer RNA (tRNA) at A site. This A1492:G530 W:W Trans base pair, in cognate tRNA bound 16S rRNAs, interacts with the second codon:anticodon canonical base pair. Cognate tRNA binding at A site induces conformational changes in the A1492, A1493 and G530 residues along with movements in domains of 30S subunit ^73^. Adenines A1492 and A1493, in a dynamic equilibrium, are either tucked in within the internal loop or flipped out. Flipped out A1493 checks for canonical base pairing for first codon:anticodon base pair via type I A-minor interaction while flipped out A1492 engages in base pairing interaction with G530. In addition, they jointly check for canonical base pairing in the second codon:anticodon interaction. Hydrogen bonding interactions and steric fit form the basis for geometry based selection of cognate tRNA over near-cognate tRNAs. Aminoglycoside antibiotics disrupt the regulation of the decoding process by locking the conformation of A1492 and A1493 in flipped out state ^74,75^. Recently it has been reported that errors in translation process can also be attributed to near-cognate codon:anticodoon noncanonical base pairs assuming geometry resembling canonical base pairing due to rare base ionization or tautomerization ^76^.

## REFERENCES

[1] T. R. Cech and J. A. Steitz, Cell, 2014, 157, 77–94.

[2] J. C. Burnett and J. J. Rossi, Chem Biol, 2012, 19, 60–71.

[3] S. Kreiter, M. Diken, S. Pascolo, S. K. Nair, K. M. Thielemans and A. Geall, J Immunol Res, 2016, 2016, year.

[4] L. E. Vandivier, S. J. Anderson, S. W. Foley and B. D. Gregory, Annu Rev Plant Biol, 2016, 67, year.

[5] B. Bradford, C. Cooper, M. Tizard, T. Doran and T. Hinton, Animal Production Science, 2016.

[6] J. G. Underwood, A. V. Uzilov, S. Katzman, C. S. Onodera, J. E. Mainzer, D. H. Mathews, T. M. Lowe, S. R. Salama and D. Haussler, Nat Methods, 2010, 7, 995–1001.

[7] J. Laborde, D. Robinson, A. Srivastava, E. Klassen and J. Zhang, Nucleic Acids Res, 2013.

[8] A. I. Petrov, C. L. Zirbel and N. B. Leontis, RNA, 2013, 19, 1327–1340.

[9] J. Roll, C. L. Zirbel, B. Sweeney, A. I. Petrov and N. Leontis, Nucleic Acids Res, 2016, gkw453.

[10] T. A. Steitz, Nat Rev Mol Cell Biol, 2008, 9, 242–253.

[11] S. A. Mortimer, M. A. Kidwell and J. A. Doudna, Nat Rev Genet, 2014, 15, 469–479.

[12] G. Suresh, H. Srinivasan, S. Nanda and U. D. Priyakumar, Biochemistry, 2016, 55, 3349–3360.

[13] C. He, Nat Chem Biol, 2010, 6, 863–865.

[14] M. Kubota, C. Tran and R. C. Spitale, Nat Chem Biol, 2015, 11, 933–941.

[15] N. B. Leontis and E. Westhof, RNA, 2001, 7, 499–512.

[16] J. Stombaugh, C. L. Zirbel, E. Westhof and N. B. Leontis, Nucleic Acids Res, 2009, 37, 2294–2312.

[17] P. S. Klosterman, M. Tamura, S. R. Holbrook and S. E. Brenner, Nucleic Acids Res, 2002, 30, 392–394.

[18] S. Halder and D. Bhattacharyya, Prog Biophys Mol Biol, 2013, 113, 264–283.

[19] J. Šponer, J. E. Šponer, A. I. Petrov and N. B. Leontis, JPhys Chem B, 2010, 114, 15723–15741.

[20] J. Šponer, A. Mokdad, J. E. Šponer, N. Spackoví, J. Leszczynski and N. B. Leontis, J Mol Biol, 2003, 330, 967–978.

[21] L. Huang, J. Wang and D. M. J. Lilley, Nucleic Acids Res, 2016, gkw201.

[22] D. Klein, T. Schmeing, P. Moore and T. Steitz, EMBO J, 2001, 20, 4214–4221.

[23] L. Huang and D. M. J. Lilley, J Mol Biol, 2016, 428, 790–801.

[24] C. C. Correll and K. Swinger, RNA, 2003, 9, 355–363.

[25] L. Jaeger, E. J. Verzemnieks and C. Geary, Nucleic Acids Res, 2009, 37, 215–230.

[26] D. Sussman, J. C. Nix and C. Wilson, Nat Struct Biol, 2000, 7, 53–57.

[27] X. Ye, A. Gorin, A. D. Ellington and D. J. Patel, Nat Struct Biol, 1996, 3, 1026–1033.

[28] P. Nissen, J. A. Ippolito, N. Ban, P. B. Moore and T. A. Steitz, Proc Natl Acad Sci, 2001, 98, 4899–4903.

[29] S. S. Ray, S. Halder, S. Kaypee and D. Bhattacharyya, Front Genet, 2012, 3, 59.

[30] J. Das, S. Mukherjee, A. Mitra and D. Bhattacharyya, J Biomol Struct Dyn, 2006, 24, 149–161.

[31] S. Bhattacharya, S. Mittal, S. Panigrahi, P. Sharma, P. S P, R. Paul, S. Halder, A. Halder, D. Bhattacharyya and A. Mitra, Database (Oxford), 2015, 2015, year.

[32] P. Čech, D. Svozil and D. Hoksza, Nucleic Acids Res, 2012, gks560.

[33] P. Čech, D. Hoksza and D. Svozil, BMC Bioinformatics, 2015, 16, 253.

[34] D. Hoksza and D. Svozil, Bioinformatics, 2012, 28, 1858–1864.

[35] D. Hoksza and D. Svozil, IEEE/ACM Trans Comput Biol Bioinform, 2015, 12, 520–530.

[36] W. Humphrey, A. Dalke and K. Schulten, J Mol Graph, 1996, 14, 33–38.

[37] Schrödinger, LLC.

[38] M. Bansal, D. Bhattacharyya and B. Ravi, Comput Appl Biosci, 1995, 11, 281–287.

[39] Y. Zhao, N. E. Schultz and D. G. Truhlar, J Chem Theory Comput, 2006, 2, 364–382.

[40] R. Ditchfield, W. J. Hehre and J. A. Pople, J Chem Phys, 1971, 54, 724–728.

[41] Y. Zhao and D. G. Truhlar, J Chem Theory Comput, 2007, 3, 289–300.

[42] F. Santoro, V. Barone and R. Improta, J Comput Chem, 2008, 29, 957–964.

[43] S. G. Stepanian, M. V. Karachevtsev, A. Y. Glamazda, V. A. Karachevtsev and L. Adamowicz, J Phys Chem A, 2009, 113, 3621–3629.

[44] W. Sun, Y. Bu and Y. Wang, J Phys Chem C, 2011, 115, 3220–3228.

[45] A. Jissy, U. Ashik and A. Datta, J Phys Chem C, 2011, 115, 12530–12546.

[46] M. Dargiewicz, M. Biczysko, R. Improta and V. Barone, Phys Chem Chem Phys, 2012, 14, 8981–8989.

[47] J. Wang, J. Gu and J. Leszczynski, J Comput Chem, 2012, 33, 1587–1593.

[48] A. Halder, A. Datta, D. Bhattacharyya and A. Mitra, J Phys Chem B, 2014, 118, 6586–6596.

[49] A. Halder, S. Bhattacharya, A. Datta, D. Bhattacharyya and A. Mitra, Phys Chem Chem Phys, 2015, 17, 26249–26263.

[50] S. F. Boys and F. Bernardi, Mol Phys, 1970, 19, 553–566.

[51] L. A. Burns, M. S. Marshall and C. D. Sherrill, J Chem Theory Comput, 2014, 10, 49–57.

[52] M. Mentel and E. J. Baerends, J Chem Theory Comput, 2014, 10, 252–267.

[53] B. Brauer, M. K. Kesharwani, S. Kozuch and J. M. L. Martin, Phys Chem Chem Phys, 2016, –.

[54] J. Řezáč and P. Hobza, Chem Rev, 2016, 116, 5038–5071.

[55] M. J. Frisch, G. W. Trucks, H. B. Schlegel, G. E. Scuseria, M. A. Robb, J. R. Cheeseman, G. Scalmani, V. Barone, B. Mennucci, G. A. Petersson, H. Nakatsuji, M. Caricato, X. Li, H. P. Hratchian, A. F. Izmaylov, J. Bloino, G. Zheng, J. L. Sonnenberg, M. Hada, M. Ehara, K. Toyota, R. Fukuda, J. Hasegawa, M. Ishida, T. Nakajima, Y. Honda, O. Kitao, H. Nakai, T. Vreven, J. A. Montgomery, Jr., J. E. Peralta, F. Ogliaro, M. Bearpark, J. J. Heyd, E. Brothers, K. N. Kudin, V. N. Staroverov, R. Kobayashi, J. Normand, K. Raghavachari, A. Rendell, J. C. Burant, S. S. Iyengar, J. Tomasi, M. Cossi, N. Rega, J. M. Millam, M. Klene, J. E. Knox, J. B. Cross, V. Bakken, C. Adamo, J. Jaramillo, R. Gomperts, R. E. Stratmann, O. Yazyev, A. J. Austin, R. Cammi, C. Pomelli, J. W. Ochterski, R. L. Martin, K. Morokuma, V. G. Zakrzewski, G. A. Voth, P. Salvador, J. J. Dannenberg, S. Dapprich, A. D. Daniels, O. Farkas, J. B. Foresman, J. V. Ortiz, J. Cioslowski and D. J. Fox, Gaussian 09 Revision C.01, Gaussian Inc. Wallingford CT 2009.

[56] R. Dennington, T. Keith and J. Millam, GaussView Version 5, Semichem Inc. Shawnee Mission KS 2009.

[57] J. Šponer, P. Jurecka and P. Hobza, J Am Chem Soc, 2004, 126, 10142–10151.

[58] P. Sharma, A. Mitra, S. Sharma, H. Singh and D. Bhattacharyya, J Biomol Struct Dyn, 2008, 25, 709–732.

[59] N. B. Leontis, J. Stombaugh and E. Westhof, Nucleic Acids Res, 2002, 30, 3497–3531.

[60] M. Sarver, C. L. Zirbel, J. Stombaugh, A. Mokdad and N. B. Leontis, J Math Biol, 2008, 56, 215–252.

[61] A. Nahvi, N. Sudarsan, M. S. Ebert, X. Zou, K. L. Brown and R. R. Breaker, Chem Biol, 2002, 9, 1043.

[62] A.G. Vitreschak, D. A. Rodionov, A. A. Mironov and M. S. Gelfand, RNA, 2003, 9, 1084–1097.

[63] A. Nahvi, J. E. Barrick and R. R. Breaker, Nucleic Acids Res, 2004, 32, 143–150.

[64] J. E. Barrick and R. R. Breaker, Genome Biol, 2007, 8, R239.

[65] J. E. Johnson, F. E. Reyes, J. T. Polaski and R. T. Batey, Nature, 2012, 492, 133–137.

[66] S. E. Butcher and A. M. Pyle, Acc Chem Res, 2011, 44, 1302–1311.

[67] M. Chawla, P. Sharma, S. Halder, D. Bhattacharyya and A. Mitra, J Chem Phys B, 2011, 115, 1469–1484.

[68] A. Halder, S. Halder, D. Bhattacharyya and A. Mitra, Phys Chem Chem Phys, 2014, 16, 18383–18396.

[69] J. C. Lee, J. J. Cannone and R. R. Gutell, J Mol Biol, 2003, 325, 65–83.

[70] V. Lisi and F. Major, RNA, 2007, 13, 1537–1545.

[71] J. L. Wilcox, A. K. Ahluwalia and P. C. Bevilacqua, Acc Chem Res, 2011, 44, 1270–1279.

[72] J. E. Sokoloski, S. A. Godfrey, S. E. Dombrowski and P. C. Bevilacqua, RNA, 2011, 17, 1775–1787.

[73] J. M. Ogle, D. E. Brodersen, W. M. Clemons, M. J. Tarry, A. P. Carter and V. Ramakrishnan, Science, 2001, 292, 897–902.

[74] H. F. Noller, Biochimie, 2006, 88, 935–941.

[75] A. Lescoute and E. Westhof, Biochimie, 2006, 88, 993–999.

[76] A. Rozov, E. Westhof, M. Yusupov and G. Yusupova, Nucleic Acids Res, 2016, gkw431.

[77] S. D. Appasamy, H. Y. Hamdani, E. I. Ramlan and M. Firdaus-Raih, Nucleic Acids Res, 2016, 44, D266–D271.

[78] J. M. Ogle and V. Ramakrishnan, Annu Rev Biochem, 2005, 74, 129–177.

[79] J. H. Davis, M. Tonelli, L. G. Scott, L. Jaeger, J. R. Williamson and S. E. Butcher, J Mol Biol, 2005, 351, 371–382.

[80] J. H. Davis, T. R. Foster, M. Tonelli and S. E. Butcher, RNA, 2007, 13, 76–86.

[81] X. Zuo, J. Wang, T. R. Foster, C. D. Schwieters, D. M. Tiede, S. E. Butcher and Y.-X. Wang, J Am Chem Soc, 2008, 130, 3292–3293.

[82] S. Baba, K.-i. Takahashi, S. Noguchi, H. Takaku, Y. Koyanagi, N. Yamamoto and G. Kawai, J Biochem, 2005, 138, 583–592.

[83] R. D. Peterson and J. Feigon, J Mol Biol, 1996, 264, 863–877.

[84] J. L. Battiste, H. Mao, N. S. Rao, R. Tan, D. Muhandiram, L. E. Kay, A. D. Frankel and J. Williamson, Science, 1996, 1547–1550.

[85] R. Stefl, F. C. Oberstrass, J. L. Hood, M. Jourdan, M. Zimmermann, L. Skrisovska, C. Maris, L. Peng, C. Hofr, R. B. Emeson et al., Cell, 2010, 143, 225–237.

[86] J. L. Chen, S. D. Kennedy and D. H. Turner, Biochemistry, 2015, 54, 3269–3285.

